# Octopamine metabolically reprograms astrocytes to confer neuroprotection against α-synuclein

**DOI:** 10.1101/2022.10.04.510813

**Authors:** Andrew Shum, Sofia Zaichick, Gregory S. McElroy, Karis D’Alessandro, Michaela Novakovic, Wesley Peng, Daayun Chung, Margaret E. Flanagan, Roger Smith, Alejandro Morales, Laetitia Stumpf, Kaitlyn McGrath, Dimitri Krainc, Marc L. Mendillo, Murali Prakriya, Navdeep S. Chandel, Gabriela Caraveo

## Abstract

Octopamine is a well-established invertebrate neurotransmitter involved in fight-or-flight responses. In mammals, its function was replaced by norepinephrine. Nevertheless, it is present at trace amounts and can modulate the release of monoamine neurotransmitters by a yet unidentified mechanism. Here, through a multidisciplinary approach utilizing *in vitro* and *in vivo* models of α-synucleinopathy, we uncovered an unprecedented role for octopamine in driving the conversion from toxic to neuroprotective astrocytes in the cerebral cortex by fostering aerobic glycolysis. Physiological levels of neuron-derived octopamine act on astrocytes via a TAAR1-Orai1-Ca^2+^-calcineurin-mediated signaling pathway to stimulate lactate secretion. Lactate uptake in neurons via the MCT2-calcineurin-dependent pathway increases ATP and prevents neurodegeneration. Pathological increases of octopamine caused by α-synuclein halts lactate production in astrocytes and short-circuits the metabolic communication to neurons. Our work provides a novel function of octopamine as a modulator of astrocyte metabolism and subsequent neuroprotection with implications to α-synucleinopathies.

## Introduction

Astrocytes, the most abundant glial cells in many parts of the central nervous system^1^, play active roles in maintaining the neuronal connectome and homeostasis. They regulate synaptic transmission and neuronal processing by recycling neurotransmitters from the synaptic space ^2–3^, are essential partners for synaptogenesis, synapse function and synaptic plasticity^3,4^, play a key role in the adaptive-protective response against oxidative stress^5,6^, and participate in neuronal energy metabolism by shuttling lactate to neurons during memory and learning^7–8,9^. While astrocyte activity is necessary to maintain neuronal health and protect neurons from toxic insults, excessive astrocyte activity (also known as reactive astrocytosis), triggered by high and sustained cytosolic Ca^2+^, has been shown to have detrimental effects on neurons^10,11–12^. Although much of our current understanding has focused on defining transcriptional signatures as well as the morphological states associated with toxic astrocytes (inflammatory, fibrous morphology, A1) vs protective astrocytes (anti-inflammatory, protoplasmic morphology, A2) ^13^, recent *in vivo* studies suggest that these signatures do not provide a full accurate description ^14^. Therefore, elucidating the extracellular signaling molecules as well as the molecular signaling mechanisms which underlie the conversion of astrocytes from neuroprotective to neurotoxic might hold important clues to understand the patho-progression of neurons in neurodegenerative diseases.

Aggregation of α-synuclein (α-syn), a small lipid binding protein, plays a central role in several neurological diseases collectively known as synucleinopathies, which include Parkinson’s Disease (PD) and Dementia with Lewy Bodies (DLB)^15^. Several studies have provided strong evidence supporting a causative role of high cytosolic Ca^2+^ as a key pathological feature in PD^16–20^. We, and others, found that the highly evolutionarily conserved Ca^2+^-dependent serine/threonine phosphatase, calcineurin, is a critical determinant of α-syn toxicity in neurons^17–20^. Further, reducing calcineurin activity pharmacologically with sub-saturating doses of the calcineurin inhibitor FK506 confers neuroprotection against α-syn in models of PD and DLB ^17–20^. A hallmark of these synucleinopathies is reactive astrocytosis ^10^. Pathological levels of Ca^2+^ have been shown to be da feature of astrocytosis ^21^. Whether the neuroprotective effects of FK506 provide neuronal rescue against the toxic effects of α-syn by regulating the protective response of astrocytes is unknown.

Here, using transcriptomic and metabolomic analyses, along with functional assays in primary cortical cultures as well as in a rodent model of α-syn pathobiology, we uncovered an unprecedented role for the trace amine, octopamine, in driving the astrocytic protective response in the cerebral cortex. Physiological neuron-derived octopamine plays an instructive role in astrocytes to undergo metabolic changes conducive for lactate production and secretion. Lactate import into neurons occurs via the MCT2-calcineurin-dependent pathway to increase ATP and prevent neurodegeneration. Pathological high levels of octopamine, caused by α-syn, halts lactate production in astrocytes and short-circuits the metabolic communication to neurons. Our work provides a novel function of octopamine as a key modulator of reactive vs protective astrocytes by harnessing their metabolism towards aerobic glycolysis and hence neuroprotection with therapeutic implications to α-synucleinopathies.

## Results

### Partial inhibition of calcineurin rewires metabolic transcription and induces a neuroprotective program in neurons and astrocytes in response to α-syn

We leveraged well-established models of α-synucleinopathy to understand the contribution of calcineurin activity to astrocytosis in the cerebral cortex^17,20,22^. Primary rat cortical cultures, consisting of glutamatergic neurons and astrocytes, were transduced with a lentivirus expressing human α-syn A53T, a mutation causing an autosomal-dominant form of PD^23^ (hereafter referred to as α-syn) or empty vector as control. α-Syn expression was regulated by the Synapsin promoter to restrict α-syn buildup to neurons where it is endogenously expressed^24,25^. After transduction, cultures were treated with a previously established sub-saturating neuroprotective dose of the calcineurin specific inhibitor, FK506^17,20^, or with vehicle as control. Differences in neuron viability were assessed by counting the number of microtubule-associated protein 2 (MAP2) positive cells (Fig. 1A,B). Consistent with our previous findings, α-syn caused approximately 50% neuron loss at 5 weeks post-transduction^17,20^. While FK506 did not have an effect in control cultures, it protected neurons against α-syn toxicity (Fig. 1A,B). The neuronal rescue was also accompanied by a reduction in phosphorylated α-syn S129, a post-translationally modified form of α-syn associated with pathology^26^ (Fig. S1A,B). Importantly, neuroprotective effects of FK506 were not due to a decrease in α-syn or calcineurin expression (Fig. S1A,C,D).

**Figure 1.**
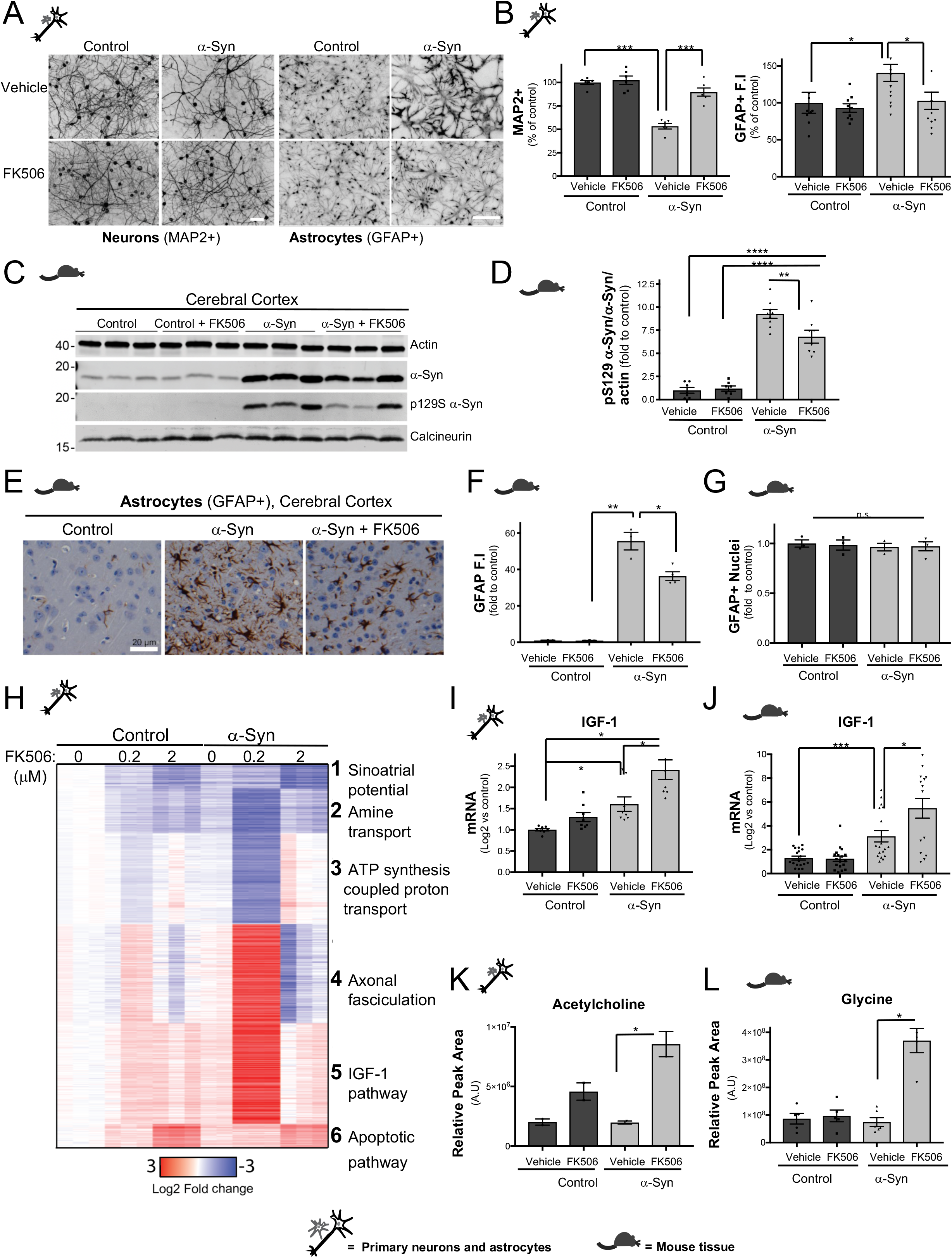
Partial inhibition of calcineurin rewires metabolic transcription and induces a neuroprotective program in neurons and astrocytes response to α-syn. **(A)** Representative immunofluorescence confocal images of neuronal (MAP2+) and astrocytes (GFAP+) from rat primary cortical cultures transduced with either control lentivirus or α-synA53T driven by the Synapsin promoter and treated with either vehicle (DMSO) or subsaturating (0.2 μM) doses of FK506 and processed for 5-6 weeks post transduction. Scale bars are 40 μm. **(B)** Quantitation of number of MAP2+ for neurons and GFAP+ fluorescence intensity (F.I) for astrocytes normalized to control-DMSO treated conditions from (A). N=9; *p< 0.05, ***p< 0.001; one-way ANOVA, *post hoc* Tukey test. **(C)** Representative Western blot for α-syn, phosphoserine-129 α-syn, and calcineurin from cerebral cortex lysates of CaMKII-Cre (control) and CaMKII-Cre-α-syn (α-syn) animals injected twice with FK506 or vehicle (DMSO) four days apart and sacrificed at day six. Actin serves as loading control. **(D)** Densitometry quantitation from the WB in (C) of p-S129 α-syn/α-syn/actin. Data is normalized to control cultures treated with vehicle (DMSO). N=3; ** p<0.01; ****p<0.0001; one-way ANOVA, *post hoc* Tukey test. **(E)** Representative immunohistochemistry for GFAP of matched sections from cerebral cortex of animals in (C). Scale bar is 20 μm. **(F)** Quantification of GFAP fluorescence intensity (F.I) from animals in (E). N≥5; * p<0.05; one-way ANOVA, *post hoc* Tukey test. **(G)** Quantification of number of GFAP positive nuclei from animals in (C). N≥5; n.s =non-significant; one-way ANOVA, *post hoc* Tukey test. **(H)** Heat map of differentially expressed genes by RNA-seq from rat primary neuronal cultures infected with either control lentivirus or α-synA53T driven by the neuronal specific Synapsin promoter, sub-saturating (0.2 μM) or saturating doses (2 μM) of FK506. All data is relative to the control cultures treated with vehicle (DMSO). Group 1, FDR 3.46E-02; Group 2, FDR 4.79E-02; Group 3, FDR 3.95E-02; Group 4, FDR 4.88E-02; Group 5, FDR 2.07E-02; Group 6, FDR 4.26E-02. **(I)** qPCR for IGF-1 normalized to control (DMSO) from cultures in (A). N≥5; *p<0.05; one-way ANOVA, *post hoc* Tukey test. **(J)** IGF-1 qPCR from cerebral cortex in (C). N≥5; * p<0.05 or ***p<0.001; one-way ANOVA, *post hoc* Tukey test. **(K)** Relative metabolite levels from cultures in (A) of acetylcholine. N=2; *p<0.05; one-way ANOVA, *post hoc* Tukey test. **(L)** Relative metabolite levels from Ctrl and α-syn transgenic mice treated with vehicle (DMSO) or FK506 from (C) of glycine. N≥5; * p<0.05; one-way ANOVA-*post hoc* Tukey test.

We next asked if expression of α-syn in neurons can cause reactive astrogliosis, a hallmark of neuronal stress/damage, previously unexplored in this model. Expression of α-syn in neurons lead to an increase in glial fibrillary acidic protein (GFAP) expression, a marker for reactive astrocytes (Fig. 1A,B). However, treatment with FK506 decreased GFAP expression without affecting the number of astrocytes (Fig. 1A,B, S1E). Importantly, we detected no differences in α-syn concentrations in the supernatants that would suggest an uptake of α-syn by astrocytes from the media (Fig.S1F).

To investigate whether calcineurin activation also played a role in astrocyte reactivity *in vivo*, we used a mouse model of α-syn pathobiology. In this synucleinopathy model, human α-syn A53T was driven by the Ca^2+^/calmodulin-dependent kinase II (CaMKII)–tTA promoter and is therefore highly expressed in the cerebral cortex, a brain region highly affected in DLB^22^. Control and α-syn expressing animals were treated with vehicle or two single doses of FK506 four days apart for a final brain content of 40 ng/g (2 ng/ml), a dose range below the standard saturating inhibitory calcineurin doses ^27^ (Fig. S1G). In the cerebral cortex, treatment with FK506 was accompanied by a reduction in phosphorylated α-syn S129, the post-translationally modified form of α-syn associated with pathology^26^, without alteration in the expression of α-syn or calcineurin (Fig. 1C,D and S1H-J). As expected, α-syn caused an increase in GFAP expression compared to control animals (Fig. 1E,F). However, treatment with FK506 decreased GFAP expression in α-syn transgenic animals without affecting the total number of astrocytes (Fig. 1E-G).

To elucidate the downstream effectors of calcineurin that convey protective responses we took a transcriptional approach using RNA sequencing (RNA-seq). Primary cortical cultures transduced with α-syn were treated with neuroprotective sub-saturating doses of FK506 (0.2μM). As controls, we utilized the non-protective saturating doses of FK506 (2μM) or vehicle alone and RNA was extracted before the onset of α-syn toxicity. In these mixed astrocyte and neuronal primary cortical cultures, α-syn expression caused a relatively modest effect on transcription, with only 233 genes being significantly differentially expressed when compared to control (Fig. 1H, S2A and Table 1). The downregulated genes were associated with pathways which are known to be impaired by α-syn, including ion transport, synaptic signaling and vesicular transport^28–30^ (Fig. S2A). The upregulated genes were associated with Nuclear factor kappa-light-chain enhancer of activated B cells (NF-KB) signaling and apoptotic processes (Fig. S2A), consistent with inflammatory and pro-apoptotic responses also previously associated with α-syn toxicity^31^. To understand the biological nature of the transcriptional changes across drug conditions, we performed a cluster analysis on the union of differentially expressed genes found in each condition compared to vehicle treated control cultures (Fig. 1H). In control cultures, saturating and sub-saturating doses of FK506 caused a similar and moderate effect relative to the vehicle control. In sharp contrast and in response to α-syn, only the neuroprotective sub-saturating doses of FK506 resulted in a striking effect on transcription, with 5164 genes being significantly differentially expressed when compared to both control and α-syn cultures. FK506 effects on α-syn cultures were specific to the neuroprotective dose since the saturating non-neuroprotective doses caused the opposite effect. Importantly, the robust transcriptional signature caused by FK506 was not due to an increase in cell number or changes in the ratios of cell types compared to control cultures treated with FK506 (Fig. S1E).

**Table 1.**
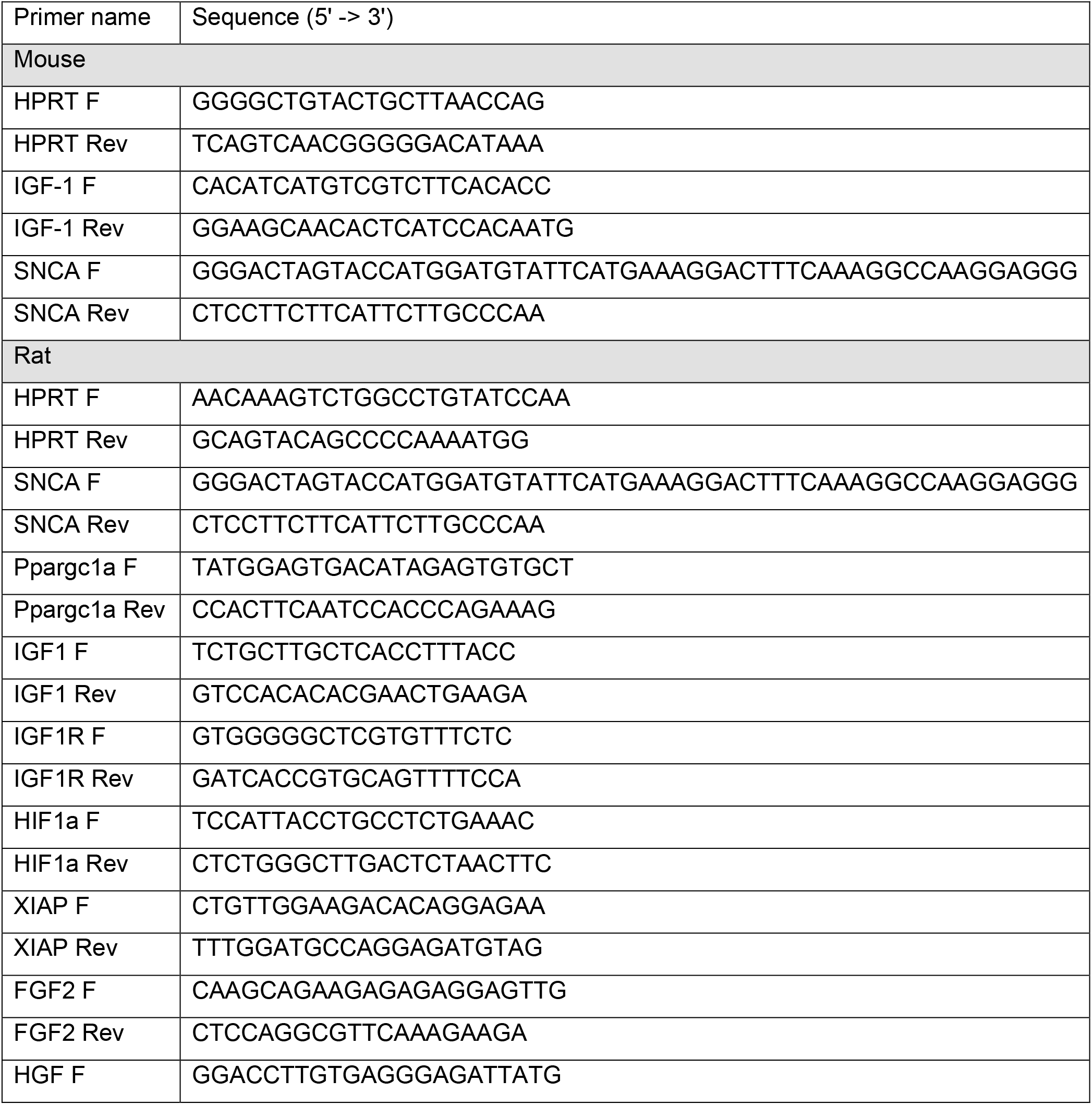
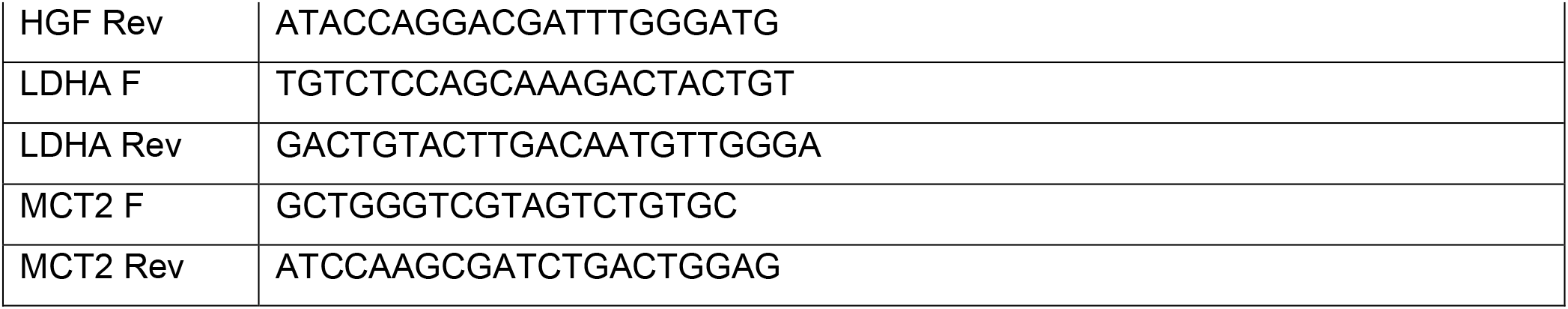
qPCR primers.

The most prominent downregulated genes in α-syn cultures treated with sub-saturating neuroprotective doses of FK506 included those belonging to amine and ATP synthesis coupled transport. In contrast, the most prominent upregulated genes corresponded to axonal fasciculation and insulin-like growth factor receptor signaling pathway. Upregulation of pro-survival and insulin-like growth factor (IGF-1) signaling pathway genes served as a proof of principle that the RNA-seq results support a protective program in response to partial inhibition of calcineurin in α-syn expressing neurons. IGF-1 signaling pathway has been shown to be neuroprotective at least partially due to inhibition of neuroinflammation^32,33^. Moreover, decreases in IGF-1 have been associated with cognitive impairment in PD and Alzheimer’s dementia ^34,35^. To validate these RNA-seq-based genes, we preformed real time PCR (qPCR) for IGF-1 in primary cortical cultures and in the cerebral cortex of α-syn transgenic animals. α-Syn caused a modest but significant transcriptional increase in IGF-1 levels compared to control, with a concordant increase in IGF-1 protein levels (Fig. 1I,J, S2J). However, treatment with FK506 in α-syn expressing neurons and transgenic animals further increased mRNA and protein IGF-1 levels (Fig. 1I,J and S2J).

Upregulation of axonal fasciculation genes also served as a proof of principle that the RNA-seq results support a neuroprotective program triggered by partial inhibition of calcineurin: neurons protected against α-syn toxicity regrew their processes (Fig.1A,B). Interestingly, the rescue in axonal fasciculation was also associated with a change in neurotransmitter profile supporting memory formation. Metabolomic analysis showed that treatment with FK506 in α-syn expressing cultures increased the levels of excitatory cholinergic-derived neurotransmitters, such as acetylcholine, and decreased the levels of inhibitory neurotransmitters such as Gamma Aminobutyric Acid (GABA) (Fig. 1K, S2K and Table 2). Treatment with FK506 in α-syn expressing mice increased the levels of glycine and decreased the elevated levels of serotonin (Fig. 1L, S2L and Table 2). Together, these data show that partial inhibition of calcineurin with FK506 transcriptionally rewires α-syn cultures towards upregulation of pro-survival pathways such as IGF-1 and axonal fasciculation and protects neurons and astrocytes against α-syn toxicity.

### Lactate promotes survival in α-syn expressing neurons in a calcineurin-dependent manner

The neuroprotective program caused by partial inhibition of calcineurin with FK506 in α-syn cultures, showed a prominent downregulation in genes belonging to ATP synthesis coupled transport. These were enriched for mitochondrial complex I subunits genes (Fig. 1H and Table 1). Downregulation of mitochondrial complex I can compromise aerobic respiration and lead to a shift towards glycolysis. We thus performed metabolomics to investigate whether a shift from oxidative phosphorylation to glycolysis was taking place in α-syn expressing cultures treated with FK506. FK506 in α-syn expressing cortical cultures caused a significant reduction in all tricarboxylic acid cycle (TCA) metabolites with a modest effect in control cultures (Fig. 2A, S3A and Table 2). Some of these metabolites were also downregulated in the cerebral cortex of FK506 treated α-syn transgenic mice (S3B and Table 2). However, one metabolite that was consistently elevated in both the primary cortical cultures and in the FK506 treated α-syn transgenic mice was lactate (Fig. 2B,C and Table 2). An increase in lactate together with a decrease in TCA metabolites suggests a shift in respiration towards glycolysis. To confirm the shift in metabolism caused by FK506 in α-syn expressing cortical cultures, we measured oxygen consumption rates in these cultures (S4A-G). Respiration measurements were not significantly changed across conditions and treatments; however, there was a trend towards an increase in the extracellular acidification rate (ECAR), a surrogate index of glycolytic flux in the α-syn expressing neurons treated with FK506 (S4G). Consistent with these trend, α-syn cortical cultures treated with FK506 expressed higher levels of hypoxia-inducible transcription factor (HIF-1α), responsible for the upregulation of glycolytic genes ^36^, by RNA-seq and qPCR (Fig. 2D and Table 1). Moreover, α-syn expressing cortical cultures and transgenic mice treated with FK506 expressed higher levels of lactate dehydrogenase 5 (LDH5), one of the five LDH isoenzymes critically involved in promoting glycolysis and highly expressed in astrocytes^37^ (Fig. 2E,F and Table 1).

**Figure 2.**
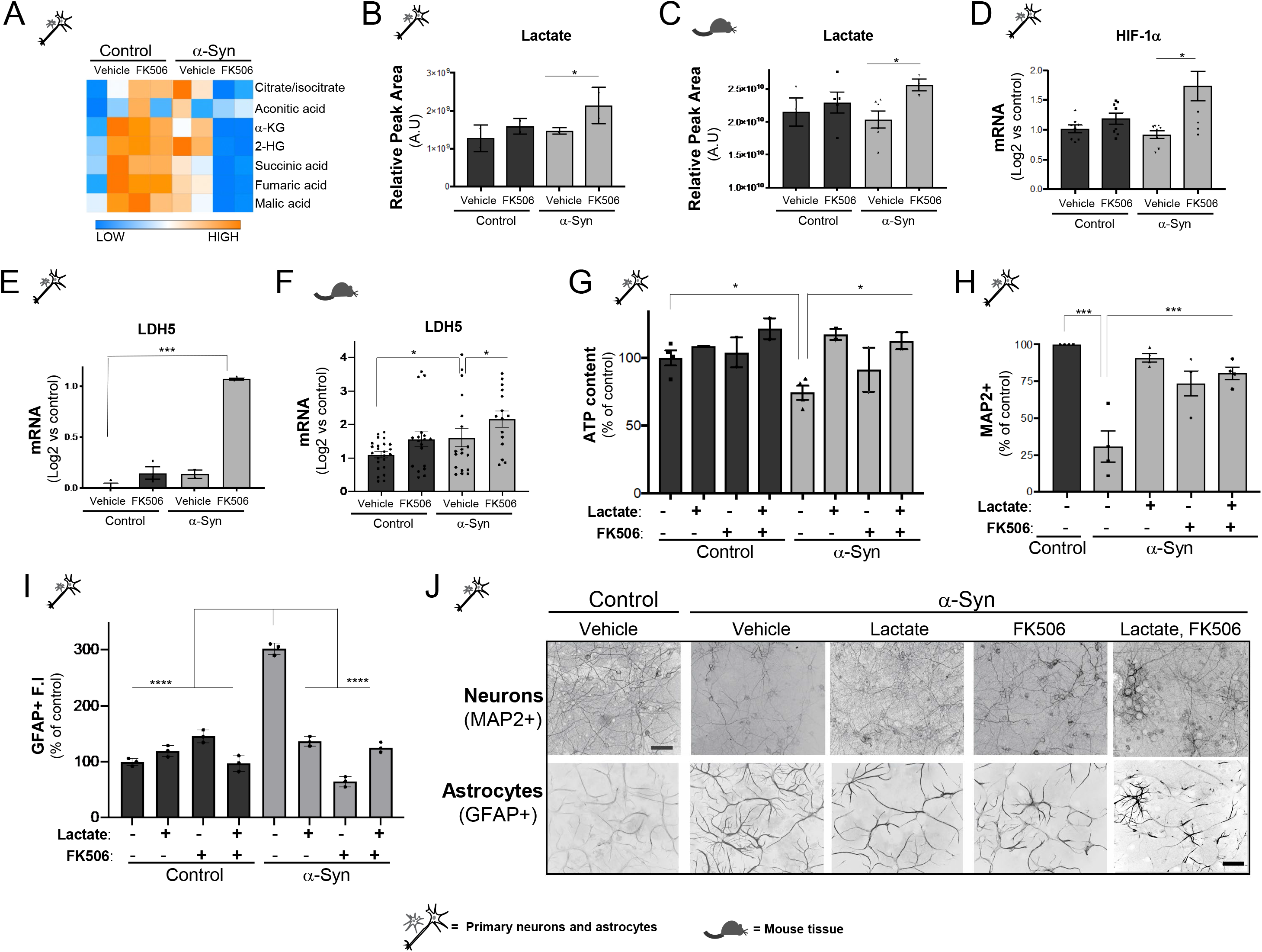
Treatment with lactate protects against α-syn toxicity in neurons and astrocytes in a calcineurin-dependent manner. **(A)** Heat map depicting the TCA cycle metabolites detected from rat primary cortical cultures transduced with either control lentivirus or α-synA53T driven by the Synapsin promoter and treated with either vehicle (DMSO) or subsaturating (0.2 μM) doses of FK506, 3 weeks post transduction (from Table 2). **(B)** Lactate metabolite from (A). N=2; *p<0.05; one-way ANOVA, *post hoc* Tukey test. **(C)** Lactate metabolite from cerebral cortex of control and α-syn transgenic animals treated with FK506, N≥5; *p<0.05; one-way ANOVA, *post hoc* Tukey test. **(D)**, qPCR for HIF-1αor LDH-5 **(E)** from primary cultures in (A). N≥3; * p<0.05. *** p<0.001; one-way ANOVA, *post hoc* Tukey test. **(F)** qPCR for LDH-5 from cerebral cortex control and α-syn transgenic animals treated with FK506. N≥5 mice, 3 technical replicates/mice; * p<0.05; one-way ANOVA with *post hoc* Tukey test. **(G)** ATP content of rat primary neuronal cultures described (A) treated with 1mM exogenous lactate, 0.2μm FK506 or both 5-6 weeks post transduction. Raw values were normalized to control-DMSO. N=6-8; *p< 0.05; one-way ANOVA, post hoc Dunnet’s multiple comparison’s test. **(H)** Number of MAP2 positive neurons in each condition normalized to control-DMSO treated conditions described in (A). N=6-8; *p< 0.05, oneway ANOVA, post hoc Tukey test. **(I)** GFAP positive fluorescence intensity (F.I) from cortical cultures in (A) normalized to control-DMSO treated conditions. N=3; 300-600 cells/biological replicate; *p< 0.001; one-way ANOVA, *post hoc* Tukey test. **(J)** Representative immunofluorescence image for MAP2+ and GFAP+ from rat primary cortical cultures in (A); scale bar = 50μm.

Aerobic glycolysis is a well-established metabolic mechanism by which cancer cells sustain rapid production of energy and metabolic intermediates that allows their survival and proliferation^38^. In the brain, astrocytes have been shown to generate lactate through aerobic glycolysis in paradigms of high energetic demands such as memory and learning ^7,39^. To test if lactate production is the mechanism by which partial inhibition of calcineurin confers neuroprotection against α-syn toxicity, control and α-syn-expressing cultures were treated with a range of lactate concentrations from nanomolar to millimolar and assayed for ATP content as a surrogate for viability. As previously established^17,20^, α-syn caused a reduction in ATP relative to control, however, treatment with 1 and 5 mM of exogenous lactate rescued these deficits (Fig.2G and S4H). Importantly, the exogenous lactate concentrations that rescued ATP are physiologic concentrations of lactate in mammals, which typically range between 1-5 mM under basal state and can reach up to 20mM during exercise^40,41^. The protective effects of 1mM lactate were also accompanied by an increase in total number of neurons as well as by a decrease in astrocyte reactivity (Fig. 2H-J and S4I-K). Importantly, the protective effects of lactate treatment were calcineurin-dependent since co-treatment with FK506 and lactate showed an epistatic relationship in α-syn expressing neurons, without any evidence of additive effect in ATP, number of neurons and astrocyte reactivity (Fig. 2G-J and S4I-K). Together, our data indicate that partial inhibition of calcineurin under α-syn promotes aerobic glycolysis and lactate-dependent neuronal survival.

### Octopamine mediates lactate production in astrocytes in a TAAR1-Orai1-Ca^2+^-calcineurin dependent manner

Neurons are mostly oxidative, whereas glial cells, namely astrocytes and oligodendrocytes, are predominantly glycolytic and hence lactate producers ^42,43^. Decreasing calcineurin activity in both, astrocyte and α-syn expressing neurons in cortical cultures promotes lactate production and survival. But how does calcineurin activity in neurons triggered by α-syn alert the neighboring astrocytes to produce lactate in a calcineurin-dependent manner? To elucidate the factor that mediates the metabolic exchange between astrocytes and neurons, we performed metabolomics on the supernatants from the mixed cortical cultures (Fig. 3A and Table 3). We detected a total of 186 metabolites across conditions with 9 being significantly different between control and α-syn expressing cultures. Of these 9 metabolites enriched for catecholamine synthesis, 3 reverted to control when treated with sub-saturating protective doses of FK506: octopamine, tryptamine and DOPET (3,4-dihydroxyphenylethanol) (Fig. 3A). However, only octopamine changed in a manner corresponding to calcineurin toxic versus protective activity: high amounts under toxic levels of calcineurin activity and lower amounts under protective levels of calcineurin activity.

**Figure 3.**
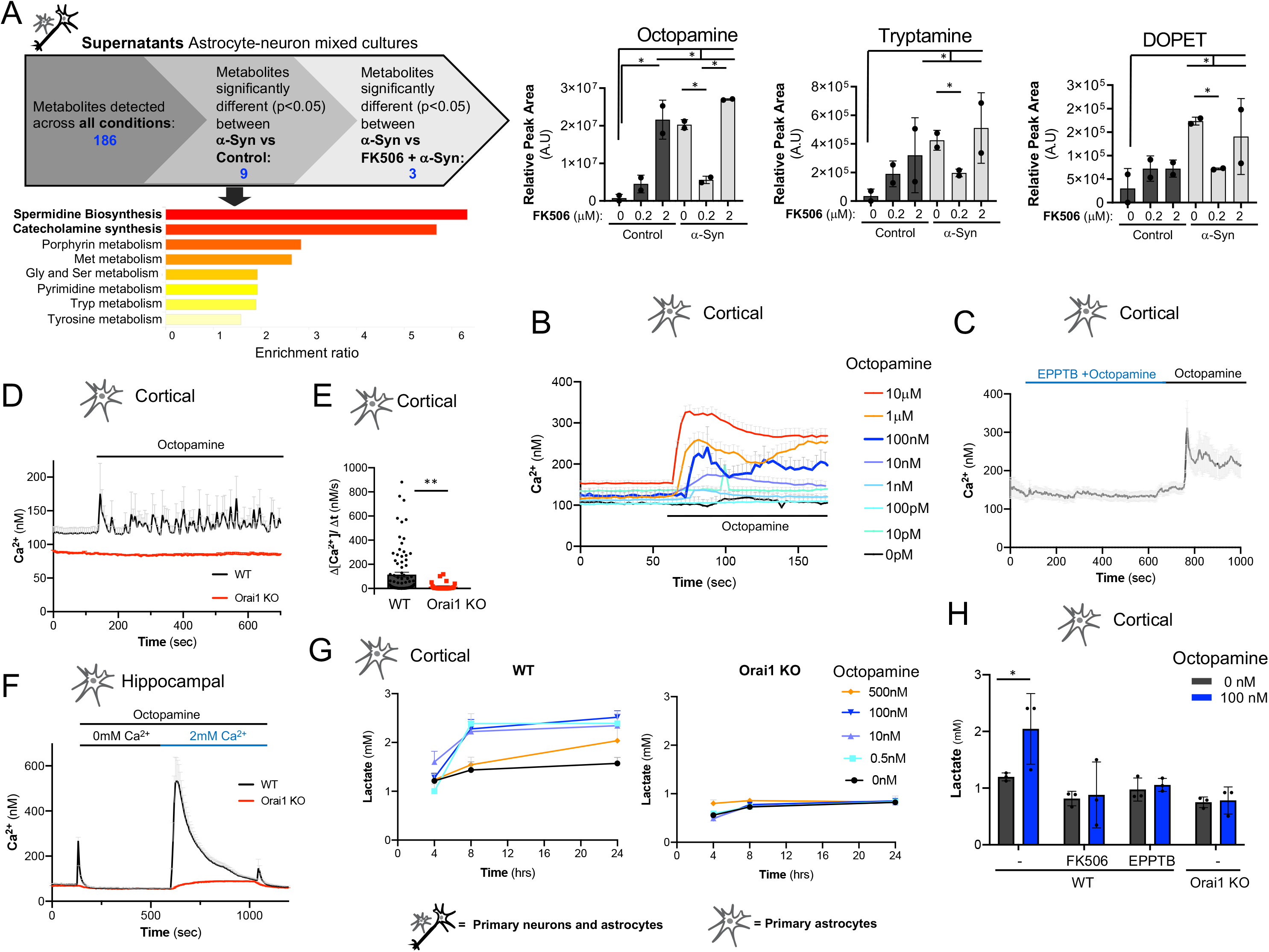
Octopamine is a key metabolite that mediates lactate production in astrocytes in a TAAR1-Orai1-Ca^2+^-calcineurin dependent manner. **(A)** Metabolite screen from supernatants of rat primary cortical cultures transduced with either control lentivirus or α-synA53T driven by the Synapsin promoter and treated with either vehicle (DMSO) or subsaturating (0.2 μM) doses of FK506 5-6 weeks post-transduction. Metabolites that were significantly different p<0.05 between control and α-syn cultures were enriched for catecholamine synthesis by Metabolite set enrichment analysis from MetaboAnalyst (McGill University). From these, only 3 hits were significantly different between α-syn and α-syn + FK506: Octopamine, Tryptamine and DOPET. N=2. *p< 0.05; one-way ANOVA, *post hoc* Dunnet’s multiple comparison’s test. **(B)** Ca^2+^ imaging of cultured rat cortical astrocytes treated with a range of octopamine concentrations (0-10μM). N=3; 7-10 cells/biological replicate. **(C)** Ca^2+^ imaging of cultured rat cortical astrocytes co-treated with octopamine (100nM) and the TAAR1 receptor inhibitor, EPPTB (500nM), washed and then challenged again with octopamine (100nM). N=3; 7-10 cells/biological replicate. **(D)** Ca^2+^ imaging of cortical astrocytes from WT or Orai1-KO mice treated with octopamine (100nM). N=3; 7-10 cells/biological replicate. **(E)** Ca^2+^ influx rates of WT and Orai1-KO cortical astrocytes from (E). N=3; 7-10 cells/biological replicate. **(F)** Ca^2+^ imaging of WT or Orai1-KO hippocampal astrocytes treated with octopamine (100nM) in the absence of extracellular calcium and in 2mM Ca^2+^. N=3; 7-10 cells/biological replicate. **(G)** Lactate measurements from WT and Orai1 KO cortical astrocyte treated with a range of octopamine concentrations (0.5-500nM) over time. N=9; *p< 0.05, one-way ANOVA, *post hoc* Tukey test. **(H)** Lactate measurements from WT and Orai1 KO cortical astrocytes pre-treated for 30 min with either FK506 (1μM) or EPPTB (200nM) and subsequently challenged with octopamine (100nM) for 4hrs. N=3. *p< 0.05; one-way ANOVA, *post hoc* Dunnet’s multiple comparison’s test.

In invertebrates, octopamine is a well-characterized neurotransmitter that plays an important role in fight-or-flight responses ^44^. Even though octopamine’s function was replaced by epinephrine during mammalian evolution, it is still present at trace amounts (nanomolar concentrations) ^45^. While octopamine does not act as a mammalian neurotransmitter given that its neuronal release is not dependent on an action potential^46^, it can still modulate neuronal function by a yet unidentified mechanism. Moreover, the importance of octopamine is highlighted by human disease where its levels are severely deregulated in PD and a range of psychiatric diseases including schizophrenia and bipolar disorder ^47,48^. To address if octopamine is the secreted neuronal metabolite responsible for modulating lactate metabolism in a Ca^2+^-dependent manner in astrocytes, we tested its ability to trigger Ca^2+^ mobilization. Pure astrocyte cultures from the cerebral cortex were loaded with the ratiometric Ca^2+^ indicator Fura-2 and challenged with a range of octopamine concentrations. While concentrations as low as 1nM where sufficient to elicit Ca^2+^ influx, the pattern of Ca^2+^ mobilization changed according to the dose. At 100nM octopamine, Ca^2+^oscillations were detected (Fig.3B and S5A). However, concentrations below 100nM triggered a monotonic low magnitude Ca^2+^ response, whereas concentrations above 100nM triggered a monotonic high magnitude Ca^2+^ response (Fig. 3B). In mammals, octopamine has been shown to bind the G-protein-coupled receptor TAAR1 (the trace amine-associated receptor 1) ^49^. While more prevalent in neurons, TAAR1 has also been shown to be expressed in astrocytes ^50^. To investigate if octopamine’s ability to mobilize Ca^2+^ was dependent on TAAR1-binding, we treated pure cortical astrocytes with the selective and reversible TAAR1 antagonist EPPTB ^51^. EPPTB abolished the Ca^2+^-dependent response of octopamine and this inhibition was reversible since it could be competed with increasing concentrations of octopamine (Fig.3C and S5B-C).

Ca^2+^ release activated Ca^2+^ (CRAC) channels, composed of Orai1 and STIM1 (stromal interaction protein 1), have been recently shown to be a major route of Ca^2+^ entry in astrocytes^52^. To investigate whether the Ca^2+^-dependent effects of octopamine were mediated by the CRAC channel, we utilized mouse Orai1 knockout (KO) cortical astrocytes, where deletion of Orai1 obliterates the channel function^53^. While WT cortical astrocytes displayed oscillatory Ca^2+^ responses in the presence of octopamine, Orai1 KO astrocytes were unaffected by octopamine (Fig. 3D,E). Importantly, octopamine’s Orai1-dependence was also observed in hippocampal astrocytes (Fig. 3F), suggesting that the Ca^2+^ mediated effects of octopamine are widespread in other regions of the brain important for cognition.

To determine if lactate synthesis is octopamine’s ultimate effector response, pure cortical astrocytes were treated with a range of octopamine concentrations in a time-dependent manner. Lysates were collected longitudinally for lactate measurements using a lactate dehydrogenase luciferase-based assay. Concentrations as low as 0.5 nm to 100nM of octopamine elicited robust lactate secretion, however concentrations of 500nM halted this effect (Fig.3G). Finally, to determine if octopamine’s effect on lactate secretion was TAAR1-Orai1-calcineurin-dependent, cortical astrocytes were co-treated with octopamine in the presence of the calcineurin inhibitor FK506, TAAR1 inhibitor, EPPTB, or in the absence of Orai1 Ca^2+^ channel. While treatment with 100nM of octopamine elicited robust lactate secretion, lactate synthesis dropped in astrocytes that were co-treated with EPPTB, FK506, or in Orai1 KO (Fig. 3H). Altogether, these data indicate that octopamine operates in a TAAR1-Orai1-calcineurin-dependent manner to regulate lactate metabolism in astrocytes in a concentration-dependent manner.

### MCT2 expression regulated by calcineurin mediates lactate neuronal survival and astrogliosis in α-syn expressing cultures

Monocarboxylase transporters (MCTs) are responsible for transporting lactate across membranes. RNA-seq and qPCR data from the primary cortical cultures showed that α-syn caused a decrease in expression of the neuronal selective MCT2^54–55^ when compared to control (Fig. 4A and Table 1). To investigate whether MCT2 expression was altered in human disease, we examined MCT2 expression in fixed postmortem tissue from humans diagnosed with two common synucleinopathies with prominent pathology in the cerebral cortex and cognitive deficits: DLB and concurrent PD and DLB. Superior frontal cortex from four to five disease cases and age-matched non-diseased controls were immunostained for MCT2 (Fig. 4B and S6). While most controls exhibited high MCT2 staining in the superior frontal cortex, MCT2 immunoreactivity was decreased in DLB and PD/DLB cases, consistent with a downregulation of MCT2 expression.

**Figure 4.**
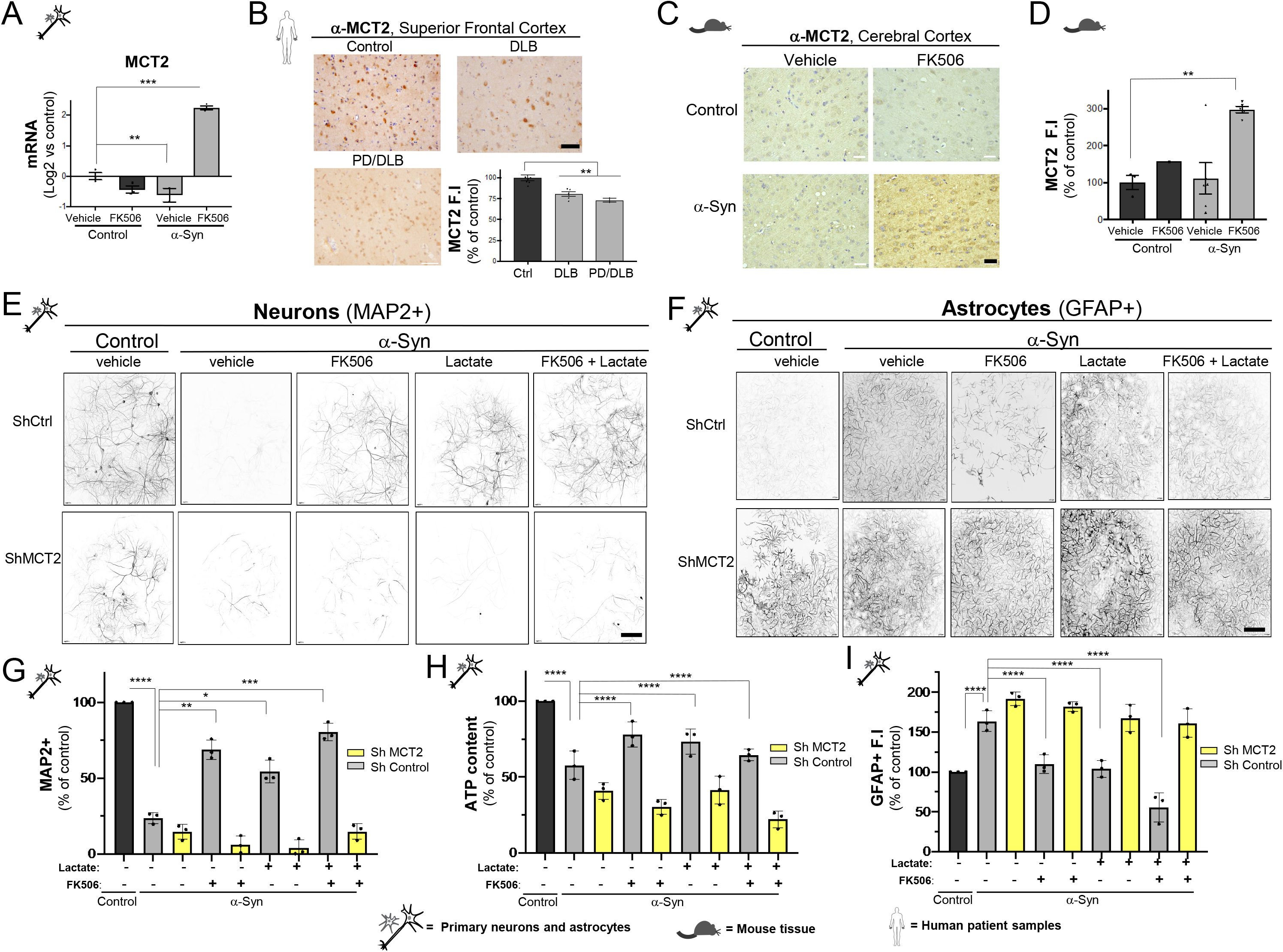
MCT2 expression regulated by calcineurin mediates lactate neuronal survival and astrogliosis in α-syn expressing cultures. **(A)** Rat primary cortical cultures transduced with either control lentivirus or α-synA53T driven by the Synapsin promoter and treated with either vehicle (DMSO) or subsaturating doses of FK506 (0.2 μM) and processed for 5-6 weeks post transduction and assayed for MCT2 by qPCR. N=3; ***p<0.001; one-way ANOVA, *post hoc* Dunnett’s multiple comparison’s test. **(B)** Representative immunohistochemistry for MCT2 in superior frontal cortex area in human DLB and PD/DLB cases and age-matched controls. An average of 5 sections from 5 control and 6 disease cases were analyzed. Scale bar is 50 μm (40x). MCT2 intensity was measured per cell. N=5-6; **p<0.01; one-way ANOVA, *post hoc* Dunnett’s multiple comparison’s test. **(C)** Representative immunohistochemistry for MCT2 from cerebral cortex of matched sections of CaMKII-Cre (control) and CaMKII-Cre-α-syn (α-syn) animals injected twice with FK506 or vehicle (DMSO) four days apart and sacrificed at day six. Scale bar is 50 μm (40x). **(D)** Quantification of average MCT2 fluorescence intensity (F.I) per cell from (C). N≥5; **p<0.01; one-way ANOVA, *post hoc* Tukey test. Data were normalized to controlvehicle (DMSO). **(E-F)** Representative immunofluorescence for MAP2+ **(E)** and for GFAP+ **(F)** from rat primary neuronal cultures co-transduced with either control lentivirus or α-synA53T driven by the Synapsin promoter and a doxycycline inducible shRNA for MCT2 or shRNA scrambled sequence as control. All cultures were treated with either vehicle (DMSO), subsaturating doses of FK506 (0.2 μM), lactate (1mM) or combinations of both lactate and FK506 once a week for 5 weeks post transduction. Scale bar = 50μM. These cultures were quantified for number of MAP2+ cells for neurons **(G)**, ATP content **(H)** and GFAP fluoresce intensity (F.I) **(I)**. Data was normalized to control-ShCtrl-DMSO condition. N=3; * p<0.05; ** p<0.01; ****p<0.0001; one-way ANOVA, *post hoc* Dunnet’s multiple comparison’s test.

Treatment with FK506 in both α-syn expressing primary cortical cultures and in α-syn transgenic mice, increased MCT2 levels by qPCR and immunostaining (Fig. 4C,D, S7 and Table 1). To determine whether regulation of MCT2 by calcineurin activity is the mechanism underlying lactate uptake in neurons to protect against α-syn toxicity, we took a knock down (KD) approach. Control and α-syn expressing neurons were co-transduced with an inducible shRNA targeting rat MCT2 or scrambled sequence as control (50% reduction levels of rat MCT2, Fig. S8A), and treated with exogenous lactate, FK506 or combinations of both. Reduction of MCT2 in control cultures had no significant effect on neuronal viability assayed by ATP content and number of remaining neurons as well as astrocyte reactivity after treatments (S8B-E). However, reduction of MCT2 in α-syn expressing neurons abolished the protective effects of FK506, lactate or the combination of both (Fig. 4E,G,H). Notably, the lack of neuronal rescue of lactate and FK506 due to reduction of MCT2 in α-syn expressing neurons was accompanied by an increase in astrocyte reactivity (Fig. 4F,I). Together, our results indicate that calcineurin regulates an MCT2-dependent lactate shuttle between astrocytes and neurons responsible for the neuroprotective effects against α-syn toxicity.

## Discussion

We had previously demonstrated the therapeutic potential of decreasing calcineurin activity in a large pre-clinical study in a rat PD model ^20^. However, the mechanism by which the downstream effectors of calcineurin activity lead to neuroprotection in the cerebral cortex had not yet been elucidated. Here, using several systems from mixed neuronal and astrocyte primary rat cortical cultures, pure astrocyte cortical and hippocampal rodent cultures, *in vivo* murine models to patient samples, we uncovered a mechanism by which decreasing calcineurin activity with FK506 leads to neuroprotection. Importantly, we demonstrate a novel role for octopamine in the mammalian brain as a key mediator of the metabolic rewiring of astrocytes and subsequent neuroprotection. We showed that moderate calcineurin activity in α-syn expressing neurons, pharmacologically achieved with sub-saturating doses of FK506, leads to release of physiological levels of octopamine and upregulation of the MCT2 transporter (S9A). In astrocytes, octopamine binds to TAAR1 and triggers an Orai1-Ca^2+-^ calcineurin-dependent signaling pathway leading to transcriptional upregulation of LDH5 and lactate synthesis. Secreted lactate is imported into neurons via MCT2, increasing ATP and neuronal survival against α-syn proteotoxic stress (S9A). Excessive calcineurin activity caused by α-syn, generates high pathological levels of octopamine and sustained Ca^2+^-calcineurin activity in astrocytes, preventing transcriptional upregulation of LDH5 and lactate synthesis. Lack of lactate import into neurons prevents ATP generation and neuronal survival (S9B).

Astrocytes have been shown to play a critical role in PD and many other neurodegenerative diseases mainly due to their neuroinflammatory toxic properties^56^. Here, we elucidate a previously unanticipated mechanism in which astrocyte reactivity can be modulated to be protective in disease: supporting aerobic glycolysis via regulation of octopamine-Ca^2+^-calcineurin activity. This neuroprotective state was defined transcriptionally by downregulation of mitochondrial complex I genes and metabolically by an increase in lactate synthesis. Therefore, our data argues that deciphering the metabolic state of reactive astrocytes is key to determine whether they will become neuroprotective or neurotoxic.

Lactate shuttles have been shown to occur in the hippocampus as a mechanism to cope for the high energy demands during memory consolidation and learning^39^. Here, we used cellular and *in vivo* models of synucleinopathy to show that therapeutic modulation of calcineurin activity can trigger a lactate shuttle that allows cells to overcome the energetic deficits caused by α-syn proteotoxic stress. The octopamine-calcineurin-dependent lactate shuttle not only allowed survival but a rewiring in neurotransmitters; from a reduction in inhibitory neurotransmission through reduction of GABA towards an increase in excitatory neurotransmission through an increase of choline-derivatives (Fig.1K,L,S2K,L). Loss of cholinergic tone has been associated with PD and other dementia-related neurodegenerative diseases such as Alzheimer’s disease ^57,58^. In fact, the drug Rivastigmine is given to patients with dementia to improve cognition by increasing cholinergic tone^59^. Our studies suggest a novel physiological role for octopamine-calcineurin-dependent lactate shuttles during cognition and possibly in the treatment of dementia-related synucleinopathies such as PD-dementia and DLB. Moreover, our studies can have therapeutic implications. First, PD patients have been shown to have lower IGF-1 levels compared to controls ^34^, and IGF-1 has been shown to be neuroprotective in models of PD and other neurodegenerative diseases^60,61^. Given that IGF-1 levels were upregulated by partial inhibition of calcineurin with the FDA-approved drug FK506 in the cerebral cortex, IGF-1 can serve as a biomarker to follow FK506 dosing in the clinic. Likewise, plasma levels of octopamine have been shown to be altered in PD ^47^. Our data demonstrated that octopamine levels are key to trigger lactate synthesis in astrocytes, and partial inhibition of calcineurin with FK506 enables this metabolic response. Therefore, octopamine can also serve as a biomarker to follow FK506 dosing in the clinic. Finally, it is well documented that exercise is beneficial to patients with PD^62^. Physical activity can have many different effects in the body, one of which is generation of lactate^63,64^. Based on our findings, lactate treatment could be explored as a novel therapeutic route to treat PD, DLB and potentially other neurodegenerative diseases where astrocytes play an important role.

## MATERIALS AND METHODS

### Primary Cortical Cultures

Embryonic rat cortical neurons were isolated from euthanized pregnant Sprague–Dawley rats at embryonic day 18 using a modified protocol from Lesuisse and Martin ^65, 66^. Protocol was approved by Northwestern University administrative panel on laboratory animal care. Embryos were harvested by Cesarean section and cerebral cortices were isolated and dissociated with 0.25% Trypsin without EDTA (Invitrogen, 15090-046) digestion for 15 min at 37°C and trituration with 1 ml plastic tip. Poly-D-Lysine (Sigma, P-1149)-coated 96-well and 24-well plates were seeded with 4 × 10^4^ and 2 × 10^5^ cells accordingly in neurobasal medium (Invitrogen, 21103-049) supplemented with 10% heat-inactivated FBS (Gibco), 0.5 mM glutamine (Gibco), penicillin (100 IU/mL), and streptomycin (100 μg/mL) (Gibco). Before seeding, cells were counted using the Automated cell counter TC10 (Bio-Rad) and viability (90-95%) was checked with Trypan Blue Stain (0.4%, Gibco 15250-061). After 1 hour (h) incubation at 37°C, media was changes to neurobasal medium (Invitrogen, 21103-049) supplemented with B27 (Invitrogen 17504044), 0.5 mM glutamine, penicillin (100 IU/mL), and streptomycin (100 μg/mL). One-half (out of 100 μl volume for 96-well plates and 500 μl volume for 24-well plates) of the media was changed 4 h before transduction on day 5 in vitro (5DIV). FK506 (Ontario Chemicals T1011), treatment (0.2 or 2 μM) of rat cortical neuronal cultures started 4 days (d) post transduction (pt) and continued twice a week (w) until αSyn toxicity was detected (usually 6.5-7.5 weeks). As a surrogate marker of cell viability, cellular ATP content was measured two-three times a week between 5-7.5 weeks post transduction using the ViaLight Plus kit (Lonza, LT07-221) or ATPlite kit (PerkinElmer, 6016941) according to the manufacturer’s instructions.

### Cell lines

HEK293T (human embryonic kidney cells) and SH-SY5Y (human neuroblastoma) cells were cultured at 37°C and 5% CO2 in Dulbecco’s Modified Eagle Medium (DMEM) with 4.5%glucose (Corning), 10% Fetal Bovine Serum (Denville) and 1X penicillin and streptomycin (Gibco). Lipid-mediated transient transfection was done using Lipofectamine 3000 (Thermofisher; L3000015) according to the manufacturer’s protocol.

### Plasmids and viruses

Plasmids: pLV-αSyn A53T (hSynapsin) promoter was a generous gift from Aftabul Haque and Susan Lindquist (Whitehead Institute for Biomedical Research, Cambridge, MA). Control (empty vector) plasmid was generated by excising αSyn and ligating the resulted product. Packaging plasmids (pMD2.G and psPAX2) were obtained from Addgene. Viruses: Lentiviral constructs were packaged via lipid-mediated transient transfection of the expression constructs and packaging plasmids (pMD2.G and psPAX2) into HEK293T cells using a modified protocol from Park et al. ^67^. Lentiviruses were purified and concentrated using the LentiX Concentrator (Clontech, PT4421-2) according to the manufacturer’s protocol. Lentivirus titer was determined using the QuickTiter Lentivirus titer kit (lentivirus-associated HIV p24; Cell Biolabs, VPK-107) or HIV-1 p24 ELISA pair set (Sino Biological Inc, SEK11695) according to the manufacturer’s protocol. All titers were further normalized to αSyn expression in SH-SY5Y cells infected with gradual multiplicities of infection (MOI) volumes of lentivirus and processed for Western blot analysis 3 dpt. Equal MOIs were used to infect rat cortical cultures on 5DIV. High levels of αSyn and MCT2 expression were confirmed by Western blot analysis, qPCR and immunofluorescence. Nontargeting ShRNAs (Horizon Discovery Dharmicon, VSC11656) or ShRNAs targeting Slc16a7 (MCT2) (Horizon Discovery Dharmicon, V3SR11254) were expressed in mammalian cells using transfection. Expression was verified visually by expression of turboRFP.

### Doxycycline treatment of cultures

Primary cortical cultures that were infected with either nontargeting ShRNAs or ShRNAs targeting Slc16a7 were treated with doxycycline. Targeted silencing of Slc16a7 (MCT2) in primary cortical cultures was induced using doxycycline treatments at a concentration of 1nM at DIV 7, 11, 14, 18 and 21 before used for experiments. Viral MOI and doxycycline titrations were performed on new batches before applied experimentally and confirmation of knockdown and expression of turbo RFP was regulated.

### Antibodies

For immunofluorescence: MAP2 (Millipore, AB5622), GFAP (Sigma, G3893) and α-synuclein (BD, 610787). Secondary antibodies used are the following: Alexa 488 (Invitrogen, a21202) and Alexa 594 (Invitrogen, a21442). For Western Blot: actin (Abcam, ab6276), α-synuclein (BD, 610787), phospho-α-synuclein (S129 (a kind gift from Dr. Takeshi Iwatsubo, The University of Tokyo, Japan), Calcineurin B (CNB, Abcam, ab154650), AKT3 (Santa Cruz, sc-134254), phospho-AKT S473 (Cell Signaling, 4060), ATM (Santa Cruz, sc-135663), Caspase-3 (Cell Signaling, 9662), p53 (Santa Cruz, sc-47698), phospho-GSK-3β S9 (Santa Cruz, sc-373800), MCT2 (Thermofisher, PA5-77498). Secondary antibodies used are the following: IRDye680 (Fisher, 925-68070) and IRDye800 (Fisher, 925-32211).

### Mice

All protocols were approved by the Massachusetts Institute of Technology university administrative panel on laboratory animal care. All live animal work was performed and concluded at MIT. However, all postmortem tissue samples were processed and analyzed at Northwestern University. C57BL6 mice with human αSyn A53T driven by the calcium/calmodulin-dependent kinase II (CaMKII)–tTA promoter were obtained from the Jackson Laboratory (Tg(tetO-SNCA*A53T)E2Cai/J, #012442) donated by Huaibin Cai’s laboratory. 12-18 old months old animals were injected with two doses of FK506 (4 mg/kg) four days apart and mice were sacrificed on day six. Blood (red blood cells were cleared out by low speed, ~1000 rpm, centrifugation) and brains were collected. Half brains were immediately flash frozen, and second halves were fixed in 4% (vol/vol) formalin. All mouse brains were analyzed for αSyn levels. Only animals with high and matched αSyn levels were picked for further analysis. At the end, six control (CAMKII^+^/αSyn^-^) injected with DMSO or FK506, six αSyn (CAMKII^+^/αSyn^+^) injected with DMSO and five αSyn (CAMKII^+^/αSyn^+^) injected with FK506 were chosen from three independent experiments that had at least five animals in each group. FK506 brain content was determined in the mouse cerebellum using liquid chromatography–MS by Sanford Burnham Prebis (SBP, La Jolla, CA). Primary astrocytes were isolated from neonatal (P0 to P3) mice as previously described (Toth et al. 2019). Briefly, hippocampus or cortex were dissected, and meninges were removed under a dissection microscope in 4°C dissection medium [l0 mM HEPES in Hanks’ balanced salt solution (HBSS)]. The tissue was digested (0.25% trypsin; Invitrogen) for l0 min in a 37°C water bath, washed twice with HBSS and dissociated gently by trituration in culture media consisting of Dulbecco’s modified Eagle’s medium with l0% fetal bovine serum and l% penicillin-streptomycin solution. Dissociated cells were filtered through a 70-μm strainer to collect cell suspension and cultured in 25-mm2 tissue culture flasks with l0 ml of medium. Half of the medium was exchanged every 3 to 4 days, and microglia were removed by forcefully shaking by hand for l0 to l5 s before each medium change. After cells reached near confluence (l2 to l4 days in vitro), medium was removed from the cells and exchanged with preheated trypsin-EDTA (0.05%). After 5 min in the incubator, culture medium was added to inactivate the trypsin, and cells were collected and centrifuged for 5 min. The supernatant was removed, and the cell pellet was resuspended in culture medium. Cells were plated on poly-L-lysine-coated glass-bottom dishes (MatTek, l4-mm diameter, l0,000 to l5,000 cells per coverslip). Plated astrocytes were maintained in the incubator and used in experiments after 2 days and within l to 2 weeks of plating. Half of the medium was exchanged with fresh medium every 4 days.

### SDS-PAGE/Western blotting

Infected neuronal cultures (5w2d post transduction) and mouse cortexes (~0.25 mg of tissue) were lysed using a radioimmunoprecipitation (RIPA) assay buffer (50mM Tris/HCl pH 7.6; 150mM NaCl; 20mM KCl; 1.5mM MgCl2; 1% NP40; 0.1% SDS). In all experiments, lysis buffer was supplemented with the Halt protease and phosphatase inhibiter cocktail (Thermofisher; 78441). Samples were incubated on ice for 30 minutes and pushed through a 27G needle (10 times) to ensure full lysis. Samples were then centrifuged at maximum RPM (~20,000 × g) for 20 minutes and the subsequent supernatants were used for Western blot analysis. Protein concentration was analyzed with the Pierce BCA Protein Assay kit (Thermofisher) and Fisherbrand™ accuSkan™ GO UV/Vis Microplate Spectrophotometer (Fisher Scientific). After the addition of the appropriate amount of the 6× Laemmli Sample Buffer (Bioland scientific LLC, sab03-02) with 5% ß-mercaptoethanol (Sigma) protein samples (10-30 μg) were boiled and separated on precasted 4-20% Criterion TGX Stain-free gels (Bio-Rad) and transferred to a nitrocellulose membrane (Amersham Protran 0.2um NC, #10600001). Membranes were blocked with 5% non-fat milk in 1X Tris-buffered saline (TBS) (50 mM Tris, pH 7.4, 150 mM NaCl) for 1 h at room temperature. Membranes were subsequently immunoblotted overnight in primary antibody at 4°C, shaking. The following day, membranes were washed three times with 1X TBST (TBS with 0.1% Tween) for 5 minutes and incubated in secondary IRDye antibody for 1 h shaking at room temperature. Membranes were washed three times with 1X TBST for before imaging using Li-Cor Odyssey® CLx Imaging System. Images were processed using Image Studio Software (LI-COR Biosciences) and signal densities were quantified using Fiji ^68^.

### IGF-1 ELISA

Cleared and normalized (to total protein) mouse cortex lysate and blood samples were used to determine levels of IGF-1 using mouse IGF-1 PicoKine ELISA kit (Boster biological technology, EK0378) and Fisherbrand™ accuSkan™ GO UV/Vis Microplate Spectrophotometer (Fisher Scientific).

### α-Syn ELISA

Neuronal cultures were seeded in poly-D-lysine pretreated 12 well plates at a density of 100,000 cells/well. After cultures were maintained for 5 and ½ weeks, media was collected, and cells were washed with 1xPBS and directly lysed with RIPA buffer containing protease inhibitors and PMSF (1mM). Lysates are scraped and blocked with 2%BSA/1xPBS before further mechanically lysed with freeze-thaw cycling. Samples are ultracentrifuged and pellets were resuspended and incubated onto 96 well ELISA plates containing 5ug/mL anti-syn42 monoclonal antibody (BD transduction #610-787). ELISAs were carried out on test samples run alongside serial diluted standard synthetic α-Syn. Samples were further incubated with α-Syn rabbit polyclonal antibody 1:500 (US biological) followed by goat anti-rabbit secondary at 1:1000 (Cell signaling) after which treated with TMB substrate. Plates were read on a microplate reader at 650nm.

### Seahorse Assays

Method was previously described (Kim, S; et. Al; 2021). OCR of neurons was analyzed using an XF 24 Extracellular Flux Analyzer (Seahorse Biosciences) according to the manufacturer’s protocol; 2.5×10^5^ neurons/well were plated on XF24 cell culture microplates. Four empty wells w/o neurons were used as background control for temperature-sensitive fluctuations in OCR analysis. Before the assay, culture medium was replaced with Seahorse XF medium (Seahorse Bioscience, #103575-100) supplemented with 1mM sodium pyruvate (Corning, #25-000-CI), 10 mM d-glucose (Sigma Aldrich, #G8769), and 2mM glutamine (Gibco, #25030-081) and incubated for 1h in a CO2-free incubator. OCR was measured after sequential injection of 1μM oligomycin, CCCP and Antimycin A. After the assay, neurons were lysed and subjected to BCA protein assay (Thermofisher, #23225) for normalization.

### mRNA isolation, cDNA synthesis and qPCR

Total RNA was isolated from mouse cortices (~0.25 mg per sample) and infected neuronal cultures (5w2d post transduction) using RNeasy kits (Qiagen, 73304&73404) according to the manufacturer’s specifications. cDNA was synthesized using High-capacity cDNA RT kit (Thermofisher, 4368814) from 0.5 μg of RNA according to the manufacturer’s specifications. qPCR was performed using iTaq Universal SYBR® Green Master Mix (Bio-Rad, 1725121) on a LightCycler 480 II (Roche). 10 pmole of primer mixes (see Table 1) and 10 ng of cDNA were used to amplify cDNA fragments. Results were expressed as ΔCp (fit point method, Roche) and obtained using the comparative C_T_ method also referred to as the 2^-ΔΔCT^ method (normalized to HPRT values).

Melt Curve analysis experiments were performed following a 2-step qPCR reaction protocol. The cDNA utilized in the PCR experiments was generated via the iScriptTM cDNA Synthesis Kit (Bio-Rad, Hercules California #1708890). PCR Reactions were then performed in triplicate (3n) using a Dye Based iTaq Universal SYBR® Green Supermix (Bio-Rad, Hercules California #1725121). The Thermal Cycling protocol was performed on a CFX96 Touch Real-Time PCR Detection System (Bio-Rad, Hercules California). Fold change quantification analysis was performed utilizing the 2-ΔΔCt method (Livak and Schmittgen 2001) to evaluate and compare genetic expression of amongst the LDHA mus, MCT2, and musHPRT genes across the samples. A parametric T-test was utilized to evaluate whether there was a significant statistical difference between replicate sample values.

### RNAseq

RNA samples from infected rat cortical neuronal cultures collected 5 weeks and 5 days post transduction were used to prepare Next Generation Sequencing (NGS) libraries. Briefly, libraries were constructed with the Illumina TruSeq Stranded mRNA Library Preparation Kit and sequenced on an Illumina NextSeq 500 at the Northwestern University, NUSeq core facility. DNA read quality control and RNA-Seq analysis was performed as described previously ^75^ with the following modifications. The quality of DNA reads, in fastq format, was evaluated using FastQC. Adapters were trimmed, and reads of poor quality or aligning to rRNA sequences were filtered. The cleaned reads were aligned to the rat genome assembly rn6 using Tophat. Gene counts were quantified from BAM files using htseq-counts. Normalization and differential gene expression were determined using EdgeR. The cutoff for determining significantly differentially expressed genes were those which exhibited a minimum two-fold change in gene expression versus the control DMSO condition and an FDR-adjusted p-value less than 0.05. Enriched gene ontology (GO) terms and P values were determined with GSEA software in the Molecular Signatures Database ^76^.

### Immunohistochemistry and Immunofluorescence

Immunohistochemistry (IHC): Histology services were provided by the Northwestern University Mouse Histology and Phenotyping Laboratory Core. Fixed half brains processed into paraffin-embedded blocks, cut and matched slices from the whole group of mice were processed for IHC staining. Initial evaluation of antibodies was tested on separate group of mice that expressed low levels of α-syn. All mounted slides were paraffin removed and rehydrated using a Leica AutostainerXL (Leica). Antigen retrieval was done in a Sodium Citrate Buffer (0.1 M, pH 6.0) for 5 min (GFAP) or 10 min (Iba1) at 95°C using a Decloaking Chamber™ NxGen (Biocare). Alternately, antigen retrieval was done for 20 min (CD3) and 5 min (CD24) at 110°C. Primary antibodies: GFAP (Biocare, #CP040A, 1:250); Iba1 (Waco, 019-19741, 1:250); MCT2 (Thermofisher, PA5-77498). Secondary antibody: biotin-SP (long spacer) AffiniPure Donkey Anti-Rabbit IgG (H+L) (Jackson immuno #711-065-152). IHC detection method: the standard avidin–biotin complex (ABC) and DAB (3,3′-diaminobenzidine) HRP substrate protocol. Images were acquired with Olympus BX41 Dual Head Microscope equipped with X-cite 120 LED camera and operated with cellSens Imaging Software (Version 1.12) provided by the core. Images were analyzed using Immunohistochemistry (IHC) Image Analysis Toolbox (FIGI) ^68^. Total GFAP and Iba1 signals were calculated using the threshold method and expressed as normalized percentage of area. Resulting data was processed using Microsoft Excel and Prizm GraphPad (http://www.graphpad.com).

### Immunofluorescence (IF)

Infected neuronal cultures (7w2d post transduction, 96-well plates) were fixed with 3% (vol/vol) paraformaldehyde. Cells were then washed three times with 1X PBS for 5 minutes each wash and permeabilized using in 0.1% Triton X-100 in 1X PBS (PBST), at room temperature (RT) for 20min and then blocked in 2% Bovine Serum Albumin (BSA, Sigma, SLBT8252) in 1X PBST for at least 60 min at RT. Primary antibody were diluted in the blocking buffer and applied over night at 4°C followed by 3X wash and incubation (1-2 h at RT) with secondary antibodies, also diluted in the blocking buffer. Subsequently, cells were 3X washed and incubated with Hoechst (Invitrogen, 33342) for 20 min at RT and washed 3X with 1X PBS. Cells were imaged using Leica DMI3000B microscope fitted with an QImaging QIClick CCD Camera and Leica HCX PL FLUOTAR L 20X/0.40 CORR PH1 objective. Image acquisition was done using Q-capture pro7 software and manual tracking of the equal exposure and digital gain setting between images. All image processing and analysis was done using Fiji ^68^. All experiments were done two independent times. Images were recorded from at least 4 wells and analyzed in a randomized and blinded fashion. MAP2-positive cell bodies (neurons) were manually calculated and subtracted from the total nuclei count (Hoechst). Fluorescence quantification was done using a Microplate Reader Infinite® M1000 PRO (Tecan) using appropriate excitation and emission wavelengths. Human brain samples were stained for MCT2 antibody (Bioss, bs-3995R) at 1:1000 dilution. Images were processed using ImageJ/FIJI software built-in package for color deconvolution (ABC-DAB) and processed for average intensity. Intensity values over six fields for each sample are compared and processed for statistical analysis.

### Astrocyte cultures

Methods were described in detail ^77^. Briefly, embryos were harvested by Cesarean section and cerebral cortices, or hippocampi were isolated and dissociated with 0.25% Trypsin without EDTA (Invitrogen, 15090-046) digestion for 15 min at 37°C and trituration with 1 ml plastic tip. Poly-D-Lysine (Sigma, P-1149)-coated 96-well and 24-well plates were seeded with 4 × 10^4^ and 2 × 10^5^ cells accordingly in neurobasal medium (Invitrogen, 21103-049) supplemented 5 ng/ml HBEGF (Sigma-Aldrich, #4643), 0.5 mM glutamine (Gibco), penicillin (100 IU/mL), and streptomycin (100 μg/mL) (Gibco). Cultures were maintained and half of culture media was exchanged once a week.

### Calcium imaging

Cortical neurons or astrocytes were seeded onto poly-D-lysine (Sigma)-coated 18 mm glass coverslips (Corning) or 23mm well fluoro dishes (World Precision Instruments). The intracellular free calcium concentration ([Ca2+]i) was measured using digital video microfluorimetry. The cells were loaded with Fura-2 acetoxymethyl ester (Molecular Probes, Eugene OR). (2 μM) for 30 min at room temperature and washed with a balanced salt solution (ACSF; containing 145mM NaCl, 5mM KCl, 2mM CaCl2, 1mM MgCl2, 10mM HEPES and 10mM glucose). Sucrose is supplemented to adjust osmolarity of the solution to 200 mOsm. Cells were allowed at least 30 min following washing to allow for dye deesterification. The glass coverslips were then mounted in a custom-designed sample chamber and perfused with ACSF solution at a rate of 1.5 ml/min by a syringe-fed system. Octopamine (Sigma-Aldrich) and EPPTB (Millipore) were applied by complete bath exchange using the perfusion system. Free Ca2+ concentration was measured by digital video microfluorimetry using an intensified CCD camera (Hamamatsu). The camera was coupled to a microscope (Nikon Diaphot), and the data acquisition was carried out by a using the Slidebook software from Intelligent Imaging Innovations (3i). Calcium concentration was calculated using calibration values as follows: Kd=135nM, Rmin=0.1986, Rmax=6.344, Beta=16.97. Influx rates were calculated by linear fits over three points prior to apex of responses.

### Lactate-Glo Assay

Lactate-Glo assay (Promega) was performed using the standard protocol defined by the manufacturer with few modifications. Briefly, 96 well plates were seeded with 40,000 cells from primary neuron cultures. After transduction with αSyn or control virus, plates are maintained for five and a half weeks with drug treatment applications weekly. For the assay, wells are washed 3x with 1xPBS before adding appropriate drug treatments (octopamine, EPPTB, FK506, or DMSO). After 30min, media is collected and processed at 4-, 8-, and 24-hour time points. Total secreted lactate concentrations are calculated using the standard curve provided by the manufacturer’s kit.

### Metabolomics experiments

#### Cell Metabolites

Soluble metabolites were extracted from primary cortical cultures using cold methanol/water (80/20, v/v) at a ratio of approximately 700 cells/uL from 200,000-400,000 cells. Soluble metabolites were extracted directly from tissue using cold methanol/water (80/20, v/v) at approximately 1μL per 50μg of tissue. Tissue was disrupted by grinding on liquid nitrogen followed by 10 seconds by ultrasonication (Branson Sonifier 250). Protein was precipitated by incubation at −80°C. Debris were pelleted by centrifugation at 18,000×g for 15 min at 4°C. The supernatant was transferred to a new tube and evaporated to dryness using a SpeedVac concentrator (Thermo Savant). Metabolites were reconstituted in 50% acetonitrile in analytical-grade water, vortex-mixed, and centrifuged to remove debris. Samples were analyzed by Ultra-High-Performance Liquid Chromatography and High-Resolution Mass Spectrometry and Tandem Mass Spectrometry (UHPLC-MS/MS). Specifically, the system consisted of a Thermo Q-Exactive in line with an electrospray source and an Ultimate3000 (Thermo) series HPLC consisting of a binary pump, degasser, and auto-sampler outfitted with an Xbridge Amide column (Waters; dimensions of 4.6mm × 100mm and a 3.5μm particle size). Mobile phase A contained 95% (vol/vol) water, 5% (vol/vol) acetonitrile, 10mM ammonium hydroxide, 10mM ammonium acetate, pH = 9.0; and mobile phase B was 100% Acetonitrile. The gradient was as follows: 0 min, 15% A; 2.5 min, 30% A; 7 min, 43% A; 16 min, 62% A; 16.1-18 min, 75% A; 18-25 min, 15% A with a flow rate of 400μL/min. The capillary of the ESI source was set to 275°C, with sheath gas at 45 arbitrary units, auxiliary gas at 5 arbitrary units, and the spray voltage at 4.0kV. In positive/negative polarity switching mode, an m/z scan range from 70 to 850 was chosen and MS1 data was collected at a resolution of 70,000. The automatic gain control (AGC) target was set at 1×10^6^ and the maximum injection time was 200 ms. The top 5 precursor ions were subsequently fragmented, in a data-dependent manner, using the higher energy collisional dissociation (HCD) cell set to 30% normalized collision energy in MS2 at a resolution power of 17,500. Data acquisition and analysis were carried out by Xcalibur 4.1 software and Tracefinder 4.1 software, respectively (both from Thermo Fisher Scientific). The peak area for each detected metabolite was normalized by the total ion current which was determined by integration of all of the recorded peaks within the acquisition window. Data were normalized by cell number and total ion current for primary cortical cultures; and by starting mass, total protein content, and total ion current for tissue.

#### Supernatant Metabolites

Soluble metabolites were extracted from primary cortical culture media using cold acetonitrile after ~24 hours in media. 20 μL media was collected from each sample and 80 μL 100% acetonitrile was added. Protein was precipitated by incubation at −80°C followed by 1 minute of vortexing. This freeze/thaw was repeated three times. After the final freeze/thaw, debris were pelleted by centrifugation at 18,000×g for 30 min at 4°C. The supernatant was transferred to a new tube for metabolomics analysis. Samples were analyzed by Ultra-High-Performance Liquid Chromatography and High-Resolution Mass Spectrometry and Tandem Mass Spectrometry (UHPLC-MS/MS). Specifically, the system consisted of a Thermo Q-Exactive in line with an electrospray source and an Ultimate3000 (Thermo) series HPLC consisting of a binary pump, degasser, and auto-sampler outfitted with an Xbridge Amide column (Waters; dimensions of 4.6mm × 100mm and a 3.5μm particle size). Mobile phase A contained 95% (vol/vol) water, 5% (vol/vol) acetonitrile, 10mM ammonium hydroxide, 10mM ammonium acetate, pH = 9.0; and mobile phase B was 100% Acetonitrile. The gradient was as follows: 0 min, 15% A; 2.5 min, 30% A; 7 min, 43% A; 16 min, 62% A; 16.1-18 min, 75% A; 18-25 min, 15% A with a flow rate of 400μL/min. The capillary of the ESI source was set to 275°C, with sheath gas at 45 arbitrary units, auxiliary gas at 5 arbitrary units, and the spray voltage at 4.0kV. In positive/negative polarity switching mode, an m/z scan range from 70 to 850 was chosen and MS1 data was collected at a resolution of 70,000. The automatic gain control (AGC) target was set at 1×10^6^ and the maximum injection time was 200 ms. The top 5 precursor ions were subsequently fragmented, in a data-dependent manner, using the higher energy collisional dissociation (HCD) cell set to 30% normalized collision energy in MS2 at a resolution power of 17,500. Data acquisition and analysis were carried out by Xcalibur 4.1 software and Tracefinder 4.1 software, respectively (both from Thermo Fisher Scientific). The peak area for each detected metabolite was normalized by the total ion current which was determined by integration of all of the recorded peaks within the acquisition window. Data were normalized by total ion count and analyzed using MetaboAnalyst 5.0.

---

**Supplemental Figure 1.**
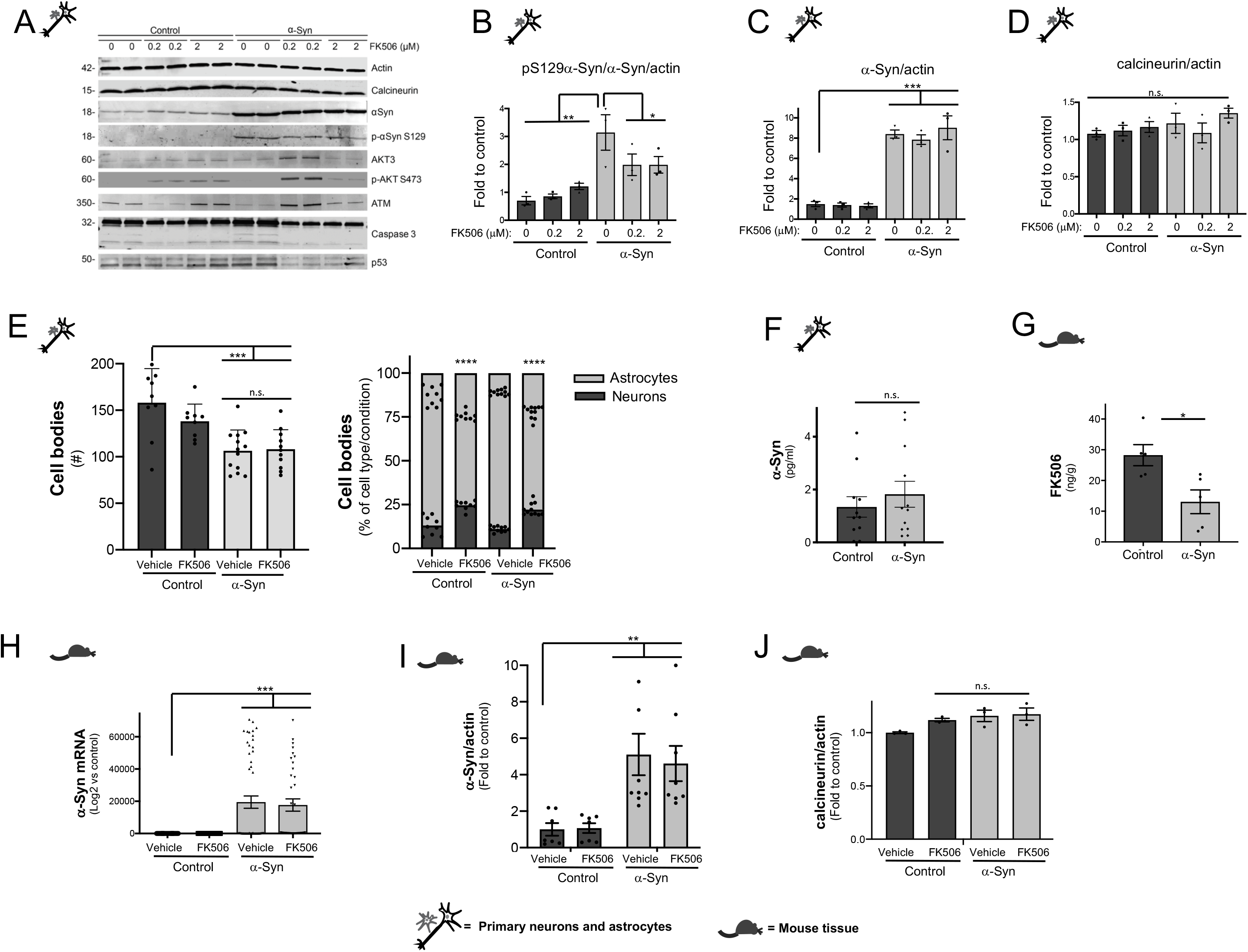
FK506 reduces pS129 α-syn *in vitro* and *in vivo*. **(A)** Representative Western blot (WB) from rat primary cortical cultures infected with either control lentivirus or α-synA53T driven by the neuronal specific Synapsin promoter, sub-saturating (0.2 μM) or saturating doses (2 μM) of FK506. **(B-D)** Densitometry quantitation from the WB in (A) of p-S129 α-syn/α-syn/actin **(B)**, α-syn/actin **(C)**, and calcineurin/actin **(D)**. All data is normalized to control cultures treated with vehicle (DMSO). N=2; *p<0.05; **p<0.01; ***p<0.001; one-way ANOVA, *post hoc* Tukey test. **(E)** Total number and percentages per condition of neurons and glia from cultures in (A). N≥6; comparison to control-vehicle; *** p<0.001, **** p<0.0001; one-way ANOVA, *post hoc* Tukey test. **(F)** α-Syn protein levels by ELISA from (A) supernatant cultures. N=9. n.s nonsignificant two-tailed t-test. **(G)** Brain FK506 levels determined by mass spectrometry from transgenic animals in Figure 1C. N≥5; * p<0.05; two-tailed t-test. **(H)** qPCR for α-syn from the indicated mice cortices. N≥5; *\*\*\** p<0.001; one-way ANOVA with *post hoc* Tukey test. **(I-J)** Densitometry quantitation from the WB in Figure 1C for α-syn/actin **(I)**, and calcineurin/actin **(J)** normalized to control-vehicle. N≥5; ** p<0.01; one-way ANOVA, *post hoc* Tukey test.

**Supplemental Figure 2.**
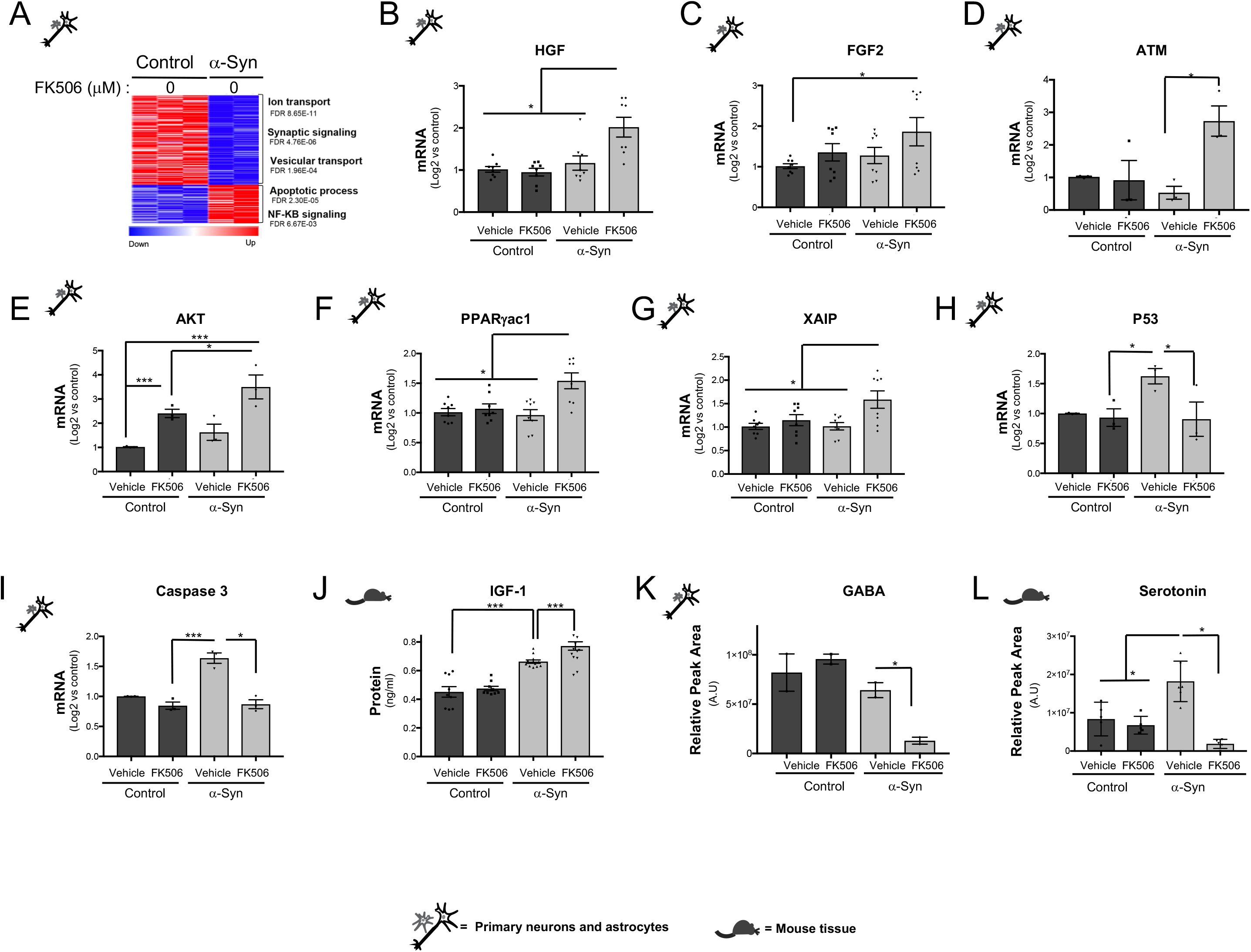
Partial inhibition of calcineurin rewires metabolic transcription and induces a neuroprotective program in neurons and astrocytes response to α-syn. **(A)** Heat map of differentially expressed genes by RNA-seq from rat primary cortical cultures infected with either control lentivirus or α-synA53T driven by the neuronal specific Synapsin promoter and treated with vehicle (DMSO) 5 weeks post-infection from table 1 and Figure 1H. **(B-I)** Series of qPCR validations from differentially expressed genes in Figure 1H for human growth factor (HGF), fibroblast growth factor (FGF-2), Adipose tissue macrophages (ATM), AKT, peroxisome proliferator-activated receptor gamma coactivator 1-alpha (PPARgac1), X-linked inhibitor of apoptosis (XAIP), tumor suppressor P53 (p53) and caspase 3. N≥5; * p<0.05; *** p<0.001; one-way ANOVA, *post hoc* Tukey test. **(J)** IGF-1 protein by ELISA from cerebral cortex of animals in Figure 1C. N≥5; *** p<0.001; one-way ANOVA with *post hoc* Tukey test. **(K)** Relative metabolite levels from cultures in (A) of GABA. N=2; *p<0.05; one-way ANOVA, *post hoc* Tukey test. **(L)** Relative metabolite levels from Ctrl and α-syn transgenic mice treated with vehicle (DMSO) or FK506 of serotonin. N≥5; * p<0.05; one-way ANOVA, *post hoc* Tukey test.

**Supplemental Figure 3.**
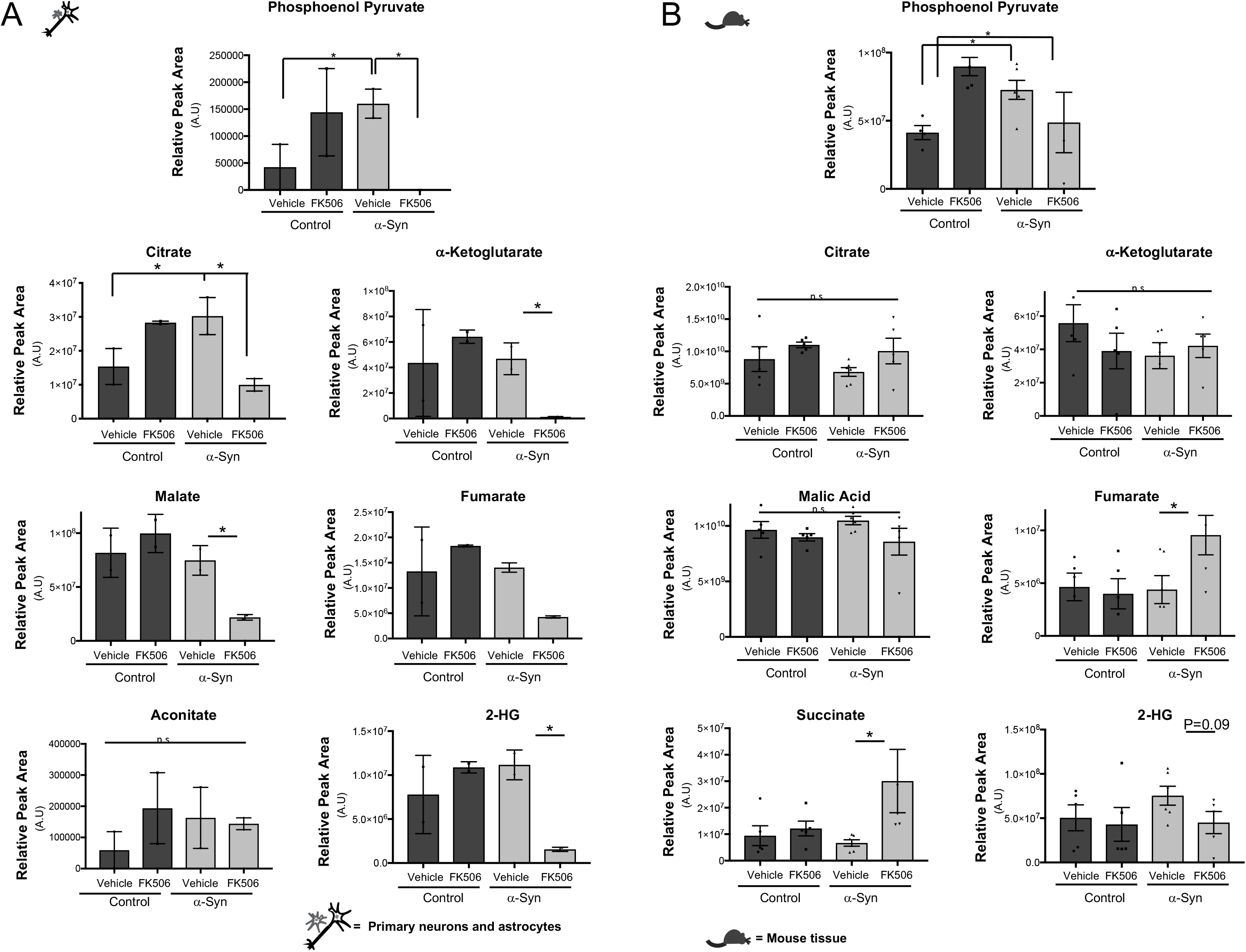
Inhibition of calcineurin in α-syn expressing neurons and astrocytes decreases TCA cycle metabolites. **(A)** Metabolites from rat primary cortical cultures transduced with either control lentivirus or α-synA53T driven by the Synapsin promoter and treated with either vehicle (DMSO) or subsaturating (0.2 μM) doses of FK506 and processed for metabolomics 3 weeks post transduction (see also Table 2). N≥2; * p<0.05; one-way ANOVA, *post hoc* Tukey test. **(B)** Metabolites from cerebral cortex of control and α-syn transgenic animals treated with FK506 or vehicle (DMSO), N≥5; * p<0.05; one-way ANOVA, *post hoc* Tukey test. Comparisons to control-vehicle, see also Table 2.

**Supplemental Figure 4.**
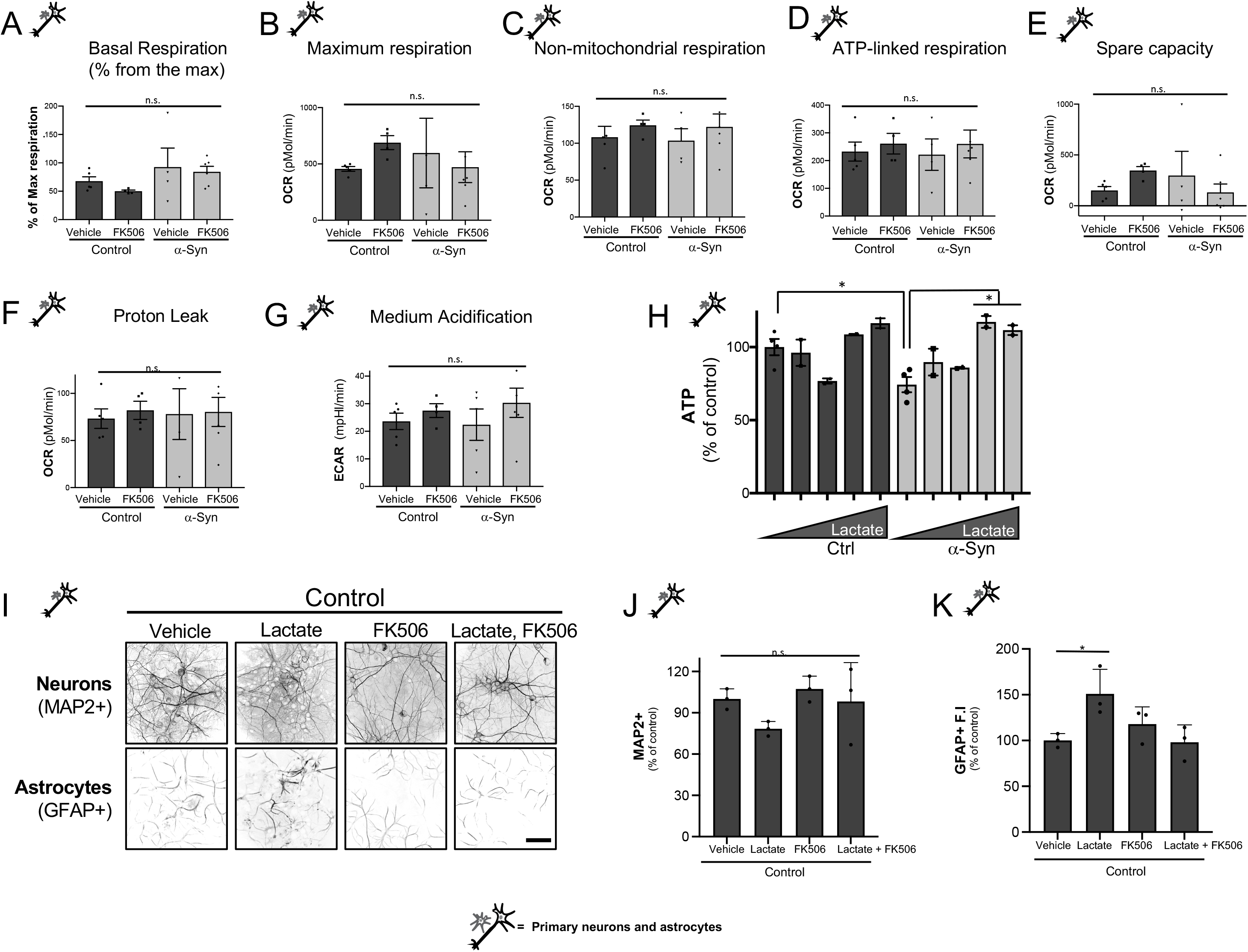
Inhibition of calcineurin in α-syn expressing cortical cultures triggers glycolysis. **(A-G)** Seahorse analysis from rat primary cortical cultures transduced with either control lentivirus or α-synA53T driven by the Synapsin promoter and treated with either vehicle (DMSO), or subsaturating (0.2 μM) doses of FK506 2-3 weeks post transduction. N=6; n.s= non-significant; one-way ANOVA, *post hoc* Dunnet’s multiple comparison’s test. **(H)** ATP content of rat primary neuronal cultures described above treated with 0, 10nM,10μM,1mM and 5mM exogenous lactate respectively for 5-6 weeks post transduction. Data were normalized to control-no lactate treatment. N=6; *p< 0.05; one-way ANOVA, *post hoc* Dunnet’s multiple comparison’s test. **(I)** Representative immunofluorescence image for MAP2+ neurons and GFAP+ astrocytes from rat primary neuronal cultures in (A) transduced with empty vector as control lentivirus; scalebar = 50μM. **(J)** Number of MAP2+ neurons normalized to control-Vehicle treated conditions described in (I). N=3; *p< 0.05, one-way ANOVA, *post hoc* Tukey test. **(K)** GFAP fluorescence intensity (F.I) normalized to control-Vehicle treated from the conditions described in (I). N=3; *p< 0.05, one-way ANOVA, *post hoc* Tukey test.

**Supplemental Figure 5.**
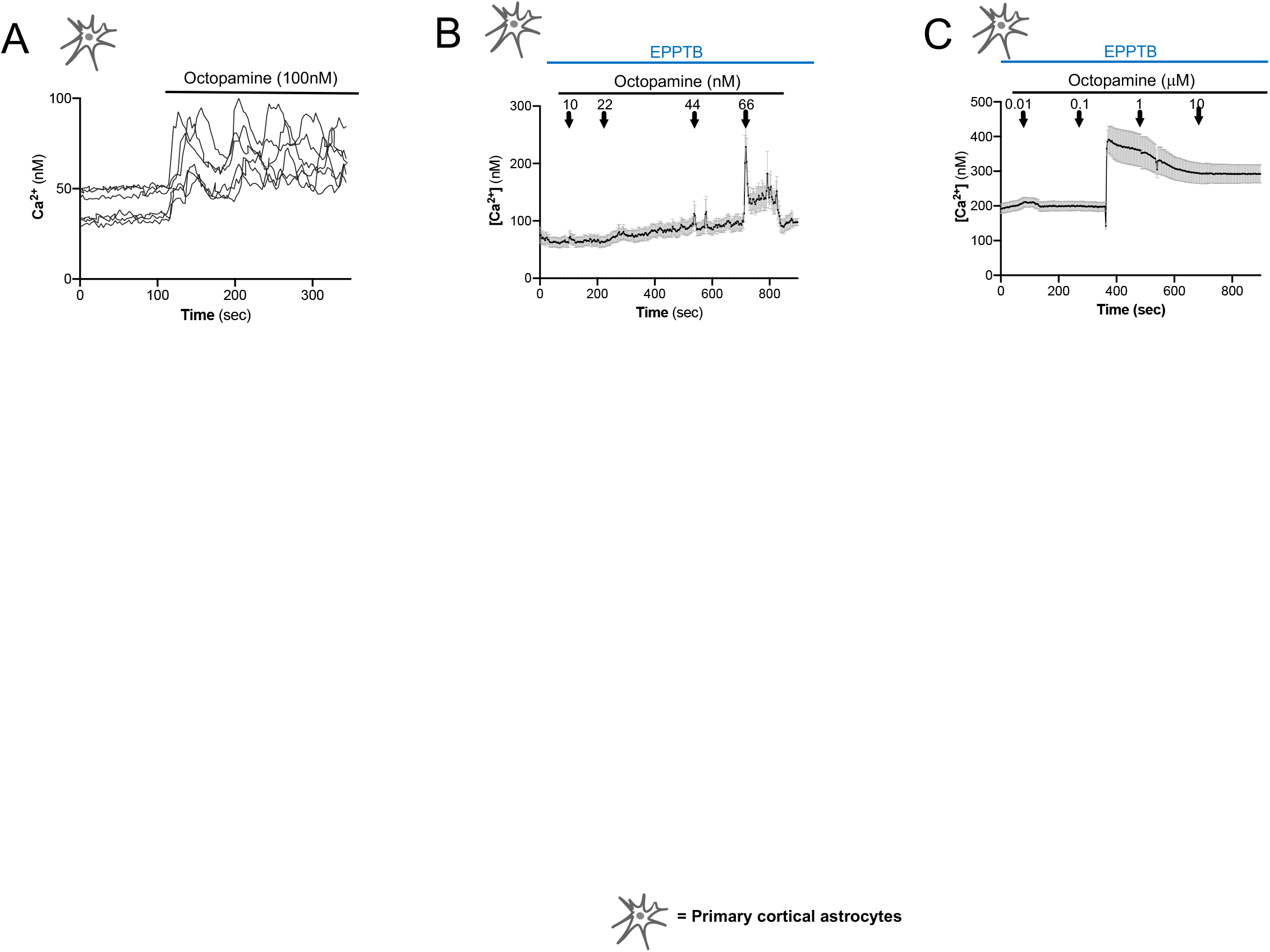
Octopamine triggers Ca^2+^ flux and lactate production in pure astrocyte cultures. **(A)** Single cell Ca^2+^ imaging traces of cultured rat cortical astrocytes treated with 100nM octopamine (N=6). **(B)** Ca^2+^ imaging traces of cultured rat cortical astrocytes treated with EPPTB (200nM) and increasing concentrations of octopamine (0-66nM). **(C)** Ca^2+^ imaging of cultured rat cortical astrocytes treated with EPPTB (200nM) and increasing concentrations of octopamine (0-10μM). All Ca^2+^ imaging is derived from N=3 biological replicates 7-10 cells/replicate.

**Supplemental Figure 6.**
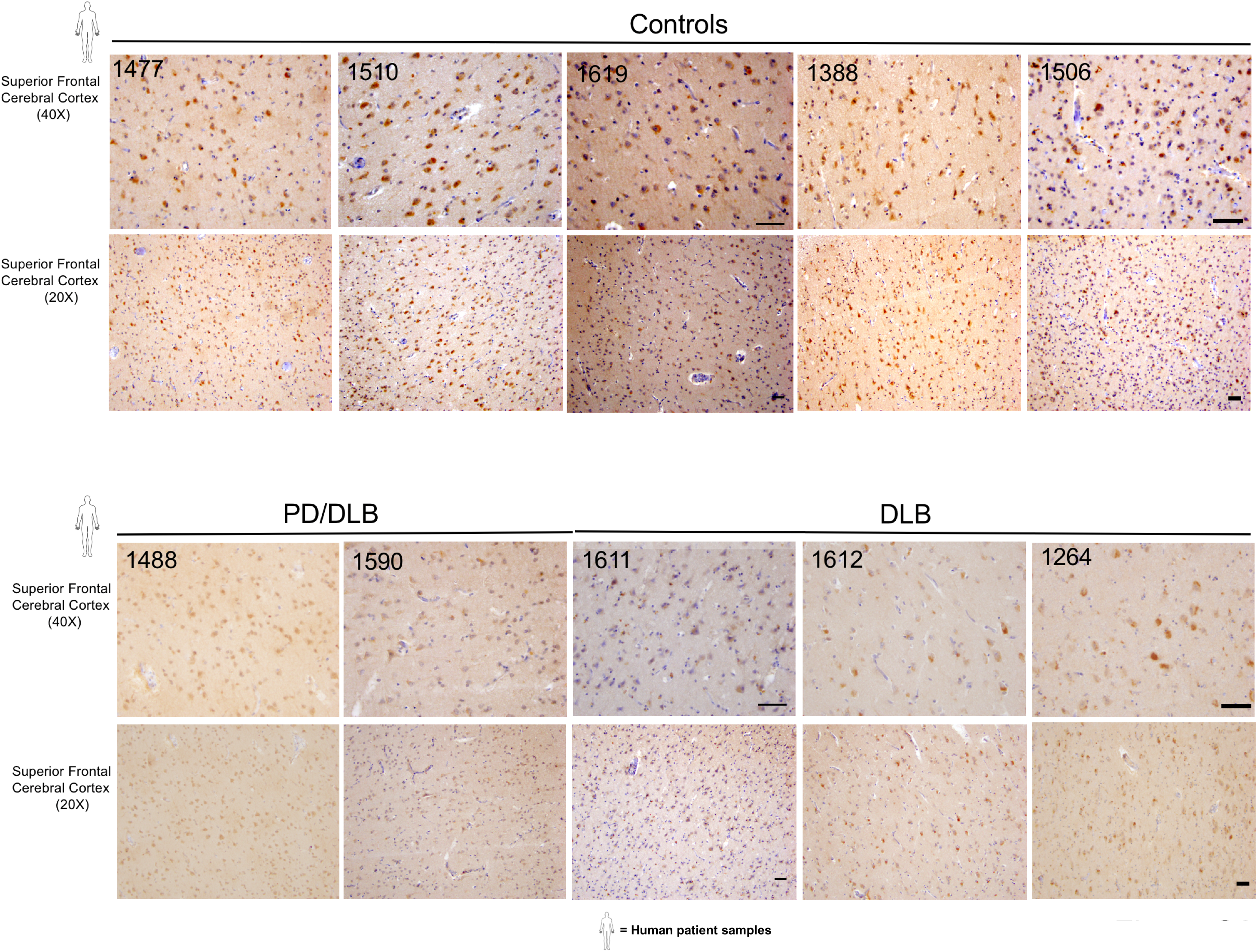
MCT2 expression is down regulated in human PD, DLB and PD/DLB disease. Individual immunohistochemistry staining for MCT2 from the superior cerebral cortex of matched sections of all control and disease cases. Scale bar is 50 μm (40×) 1000 μm (20×).

**Supplemental Figure 7.**
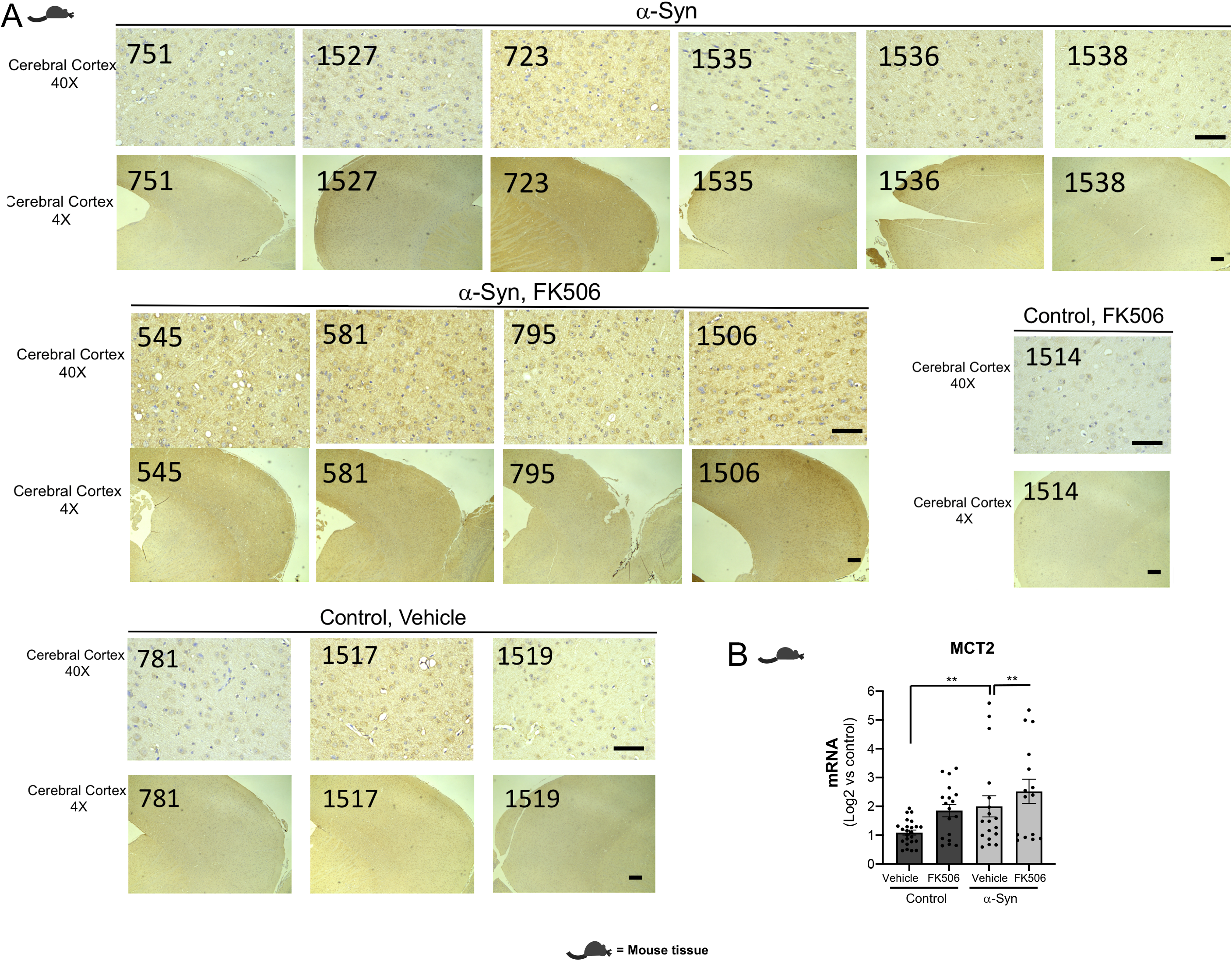
Calcineurin regulates MCT2 expression in α-syn transgenic animals. **(A)** Individual immunohistochemistry staining for MCT2 from cerebral cortex of matched sections of all control and α-syn transgenic animals treated with FK506. 50 μm(40×) 1000 μm (4x). **(B)** MCT2 expression by qPCR from animals in (A). N≥5 animals, 3 technical replicates/animal; **p<0.01; One-way ANOVA *post hoc* Dunnett’s multiple comparison’s test.

**Supplementary Figure 8.**
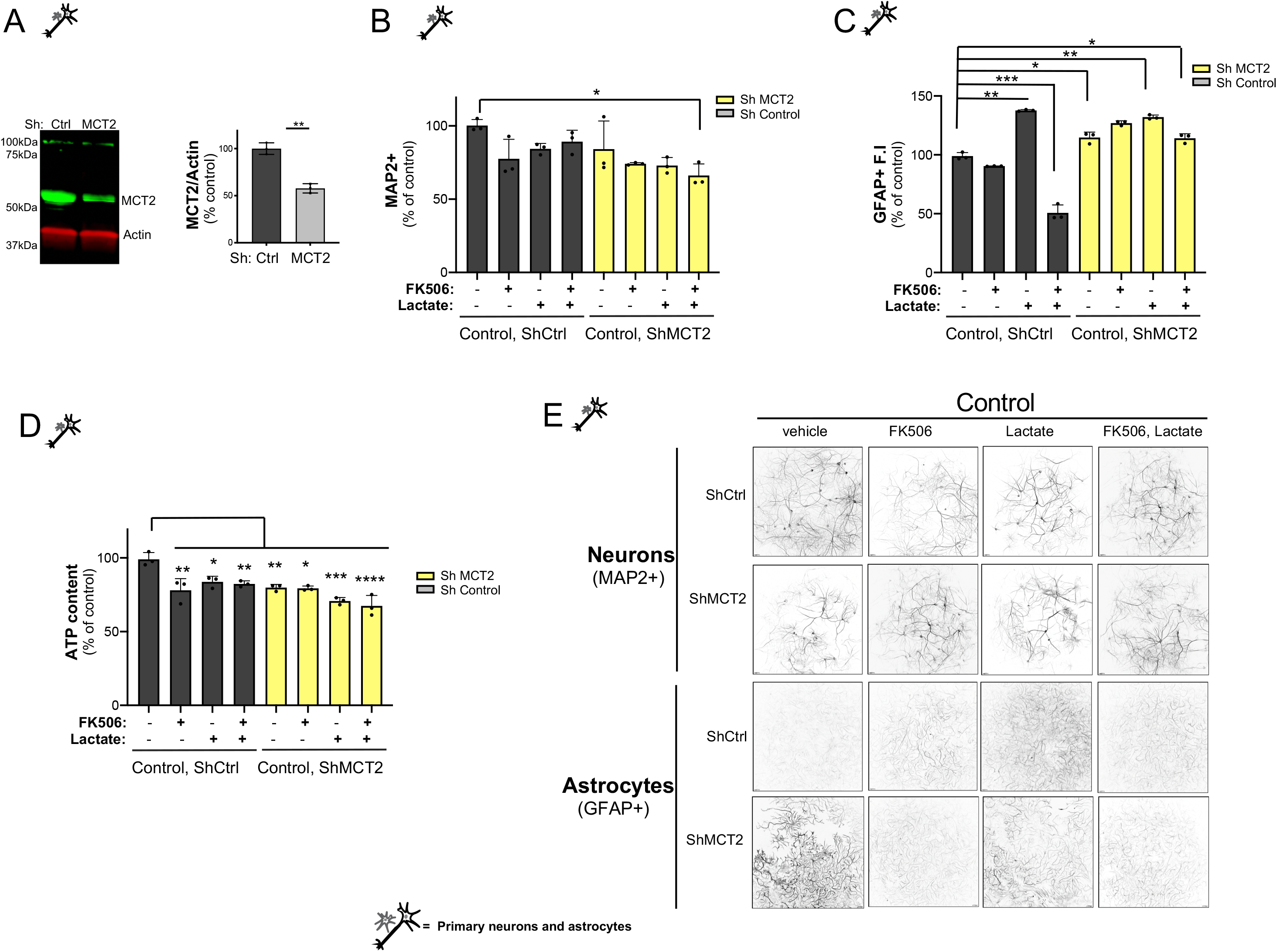
Treatment with lactate promotes survival in α-syn expressing neurons and astrocytes in a calcineurin dependent manner. **(A)** Representative WB for MCT2 expression in primary cortical cultures transduced with shRNA control or shRNA MCT2. Actin serves as a loading control. Quantitation of the WB after normalization for actin. N=2; **p<0.05; two-tailed t-test. **(B-E)** Rat primary neuronal cultures co-transduced with either control lentivirus driven by the Synapsin promoter and a doxycycline inducible shRNA for MCT2 or shRNA scrambled sequence as control. All cultures were treated with either vehicle (DMSO) or subsaturating (0.2 μM) doses of FK506, lactate (1mM) or combinations of both lactate and FK506 once a week for 5 weeks post transduction. Cultures were assayed for number of MAP2+ neurons **(B),** GFAP fluorescence intensity (F.I) **(C),** and for ATP content **(D)**. Data was normalized relative to control-ShCtrl-Vehicle. N=4-6; * p<0.05; ** p<0.01; **** p<0.0001.one-way ANOVA, *post hoc* Dunnett’s multiple comparison’s test. **(E)** Representative immunohistochemistry for MAP2+ neurons and GFAP+ astrocytes from (B-C). Scale bar = 50μm.

**Supplemental Figure 9.**
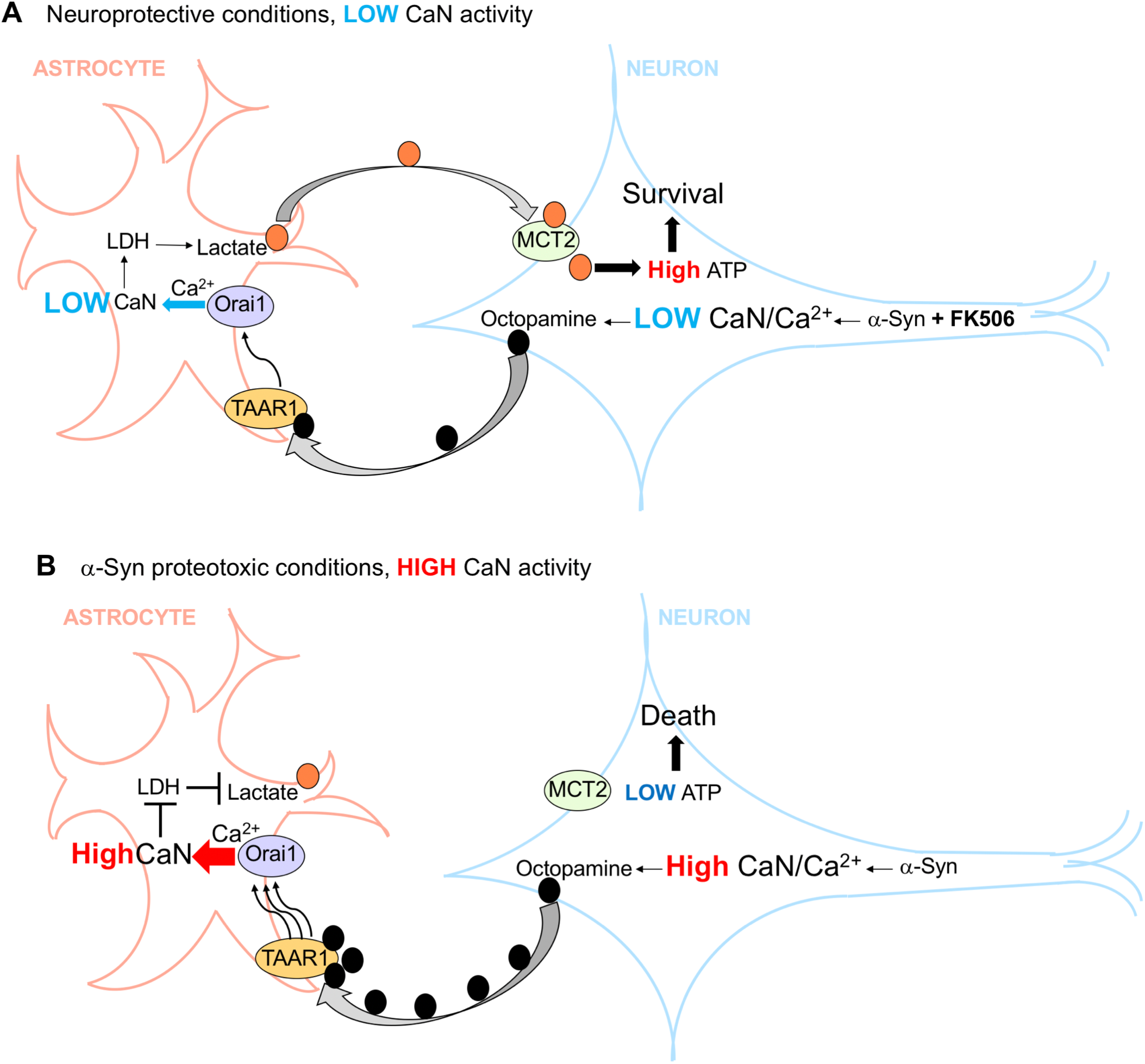
Diagram representing octopamine’s mode of action in astrocytes and neurons under α-synuclein. **(A)** Under neuroprotective conditions, low calcineurin activity in neurons, achieved with subsaturating doses of FK506, leads to release of physiological levels of octopamine which binds to the G-protein couple receptor TAAR1 to activate Orai1 in astrocytes. Oscillatory Ca^2+^ influx though Orai1 activates calcineurin, and leads to transcriptional upregulation of LDH and subsequent lactate production in astrocytes. Secreted lactate is taken up by neurons via MCT2 increasing neuronal ATP and survival. **(B)** Under neurotoxic conditions, high calcineurin activity, caused by α-syn in neurons, releases high pathologic levels of octopamine which leads to high and sustained Orai1-dependent Ca^2+^ flux in astrocytes. High calcineurin activity in astrocytes halts LDH transcription and subsequent lactate production. Lack of lactate in neurons prevents ATP generation and leads to neuronal death.

